# A Rare T-Cell Factor 4 Lineage-negative Epithelial Stem Cell Supports Wound Repair and APC-deletion-induced Colon Tumorigenesis

**DOI:** 10.64898/2026.03.17.712502

**Authors:** Annika V. Thorpe, Tim Mosbruger, Stephanie J. Georges, Olivia M. Crowley, Therese Tuohy, Brian Dalley, Benjamin D. Bice, Andrew K. Fuller, Julio R. Hidalgo, Christopher D. Green, Saher Sue Hammoud, Melinda L. Angus-Hill

## Abstract

To maintain barrier homeostasis, the colonic and intestinal epithelial lining is continually renewed by rapidly proliferating epithelial crypt base columnar (CBC) stem cells that reside at the base of crypts. Using mouse lineage tracing, immunohistochemistry, and single-cell sequencing, we have identified a rare, non-CBC, T-cell factor 4 lineage-negative (*Tcf4* Lin-) stem cell population that gives rise to secretory and absorptive precursors. Following endoscopic biopsy-induced injury, *Tcf4* Lin- stem cells are recruited to the wound bed and to the site of expanding crypts and function in barrier restoration and wound repair. We show that in a *Tcf4*-haploinsufficient background, the *Tcf4* Lin-, but not the *Tcf4* Lin+, cell population represents the cell of origin for colon tumors driven by deletion of *Apc*. Our results provide a foundation for understanding *Apc*-allele-specific differences during colon tumorigenesis and identify a new stem-cell population that may prove valuable in the treatment of diseases caused by intestinal barrier homeostasis defects.

## Introduction

The colonic and intestinal epithelium is continuously self-renewing. In the crypt, stem cells give rise to transit amplifying cells (TA cells) that proliferate, migrate and develop into terminally differentiated absorptive or secretory lineages. The intestinal stem-cell field has largely focused on the identification and characterization of stem cells in the small intestine, with relatively few studies in normal colon until recently. The most rigorously studied and extensively reviewed stem-cell marker in both small-intestine and colon has been *Lgr5*. Expression of this *Wnt* target gene marks a stem cell known as the crypt base columnar (CBC) cell (recently reviewed in Barker, 2014). Contrary to long-held notions of stem cells, these cells are long-lived and rapidly dividing (Barker et al., 2008b). A more quiescent intestinal stem-cell population is marked by expression of *Bmi1*, *Lrig1*, *Tert*, and *HopX*. Whether this quiescent stem-cell population is independent from CBC cells is controversial, since its markers are also found in rapidly dividing *Lgr5*+ stem cells (Munoz et al., 2012). In support of a distinct quiescent cell population, the *Lgr5*-expressing stem cells are dispensable in the small intestine, suggesting that non-*Lgr5+* stem cells are sufficient to maintain intestinal homeostasis (Tian et al., 2011). Furthermore, *Lgr5*- and *Lrig1*-promoter-driven conditional mutagenesis of the tumor suppressor gene *Apc* gave rise to different tumor phenotypes. *Lrig1*-promoter-driven heterozygous *Apc^LoxEx14^*deletion drove tumorigenesis efficiently, while *Lgr5*-promoter-driven *Apc^LoxEx14^* deletion required homozygosity, and the extent of tumorigenesis was not clearly defined (Barker et al., 2009; Powell et al., 2014). Thus, *Lrig1* and *Lgr5* appear to mark two distinct stem-cell populations.

The *Wnt* signaling pathway is driven by *β-catenin*, and along with T-cell factor 4 (*Tcf4*, also known as *Tcf7l2*), is thought to be important for maintenance of the intestinal stem-cell population (Korinek et al., 1998). Indeed, downregulation of the β-catenin-TCF/LEF pathway is correlated with differentiation (van de Wetering et al., 2002). However, in the colon, TCF4 protein is expressed at very low levels in proliferative cells at the base of the crypt and at much higher levels in differentiated cells further up the crypt (Angus-Hill et al., 2011). Still, *Tcf4* is thought to be important for stem-cell self-renewal, since loss of this gene results in depletion of the stem-cell compartment in the duodenum of newborn animals (Korinek et al., 1998). Due to this finding, TCF4 is thought to be an oncogenic protein that acts in a positive way to *promote* intestinal cell growth and tumorigenesis (Korinek et al., 1998). However, recent studies have challenged this long-standing paradigm, as loss-of-function mutations in *TCF4* were detected in primary human colorectal tumors and were shown to *stimulate* the growth of colon cancer cells *in vitro* (Tang et al., 2008; Wood et al., 2007).

Inactivation of both alleles of the Wnt pathway inhibitor gene Adenomatous polyposis coli (*APC*) is an early event in most colon adenomas and carcinomas (Kwong and Dove, 2009), and several mouse models have been developed that contain *Apc* mutations similar to those associated with human disease (reviewed in McIntyre et al., 2015). Conditional *Apc* alleles, in particular, have been successful in modeling the colon-specific development of tumors (Zeineldin and Neufeld, 2013). It has long been postulated that mutation of one *Apc* allele within the “mutation cluster region” results in an N-terminal truncated allele that is required for tumor initiation, suggesting either a gain-of-function or dominant-negative role for the mutant allele (Sieber et al., 2002). However, whole-gene *APC* deletion has been identified in humans with classical familial adenomatous polyposis (FAP), raising the possibility that *APC* truncation, *per se*, is not essential for tumorigenesis (Sieber et al., 2002). This possibility was further supported by driving conditional *Apc* deletion (*Apc^LoxEx1-15^*) in all intestinal epithelium using the *Villin^Cre^* driver, where mice have significantly decreased median survival and on average more colon tumors than *Apc^Min^* (truncation at amino acid 850) animals, further calling into question the supposed requirement for an *Apc* gain of function for tumorigenesis (Cheung et al., 2010).

To rigorously investigate the function of TCF4 in carcinogenesis, we generated mouse lines in which *Tcf4* protein expression is effectively abolished. Heterozygous mutation of *Tcf4* dramatically *increased* the number of colon tumors in the *Apc^Min^* model that normally only develops small intestinal tumors, suggesting a role for *Tcf4* in tumor suppression, *<u>not</u>*tumor promotion (Angus-Hill et al., 2011). Importantly, colon tumors from these mice were similar to those that develop in humans. This indicated that *Tcf4* acts as a tumor suppressor rather than an oncogene. We also discovered that *Tcf4* plays a key role as an inducer of terminal differentiation of intestinal cells, and we postulated that this feature may account, at least in part, for the tumor-suppressive function of *Tcf4* (Angus-Hill et al., 2011). Further, loss of *Tcf4* in *Tcf4*-expressing cells results in colon epithelial cell necrosis and crypt hyperplasia (Angus-Hill et al., 2011). However, using lineage analysis, we show here that these proliferative hyperplastic cells have never in fact undergone activation of the *Tcf4* promoter and that loss of *Tcf4* from cells that would normally express it must therefore promote hyperplasia in a non-cell-autonomous manner. To understand what is driving these *Tcf4*-lineage-negative (*Tcf4* Lin-) cells to undergo hyperplasia in response to loss of *Tcf4*, we have isolated and characterized them using *Tcf4*-lineage-specific mouse reporter lines. We found that the *Tcf4* Lin- cell population is comprised of both epithelial and immune cells, and that the epithelial (Epi) population can be further subdivided into a stem-cell population as well as absorptive and secretory precursors. Importantly, Epi *Tcf4* Lin- stem cells represent a very rare population of cells that appears to be distinct from, and independent of, other previously characterized intestinal stem-cell populations. Following mechanical colonic damage, Epi *Tcf4* Lin- cells are recruited to colonic lesions and proliferate to aid in crypt expansion into the wound. Using these *Tcf4*-lineage-specific reporter lines, we further show that heterozygous truncation but not deletion of *Apc*, in a *Tcf4*-haploinsufficient background, drives colon tumorigenesis in *Tcf4* Lin+ cells and that the *Tcf4* Lin- cell represents the cell of origin for tumors arising from *Apc* deletion. Thus, the nature of the *Apc* mutation dictates the colon-cell lineage from which it can promote tumorigenesis.

## Results

### *In vivo* characterization of *Tcf4* Lin- cells suggests the presence of a stem cell population that gives rise to epithelial descendants

To define the lineage of cells expressing *Tcf4* in the colon, we crossed each of two transgenic lines in which the *Tcf4* promoter drives expression of the *Cre* recombinase gene, *Tcf4^CreNeo^* and *Tcf4^Cre^*, to the *Rosa^mTmG^* mouse, a Cre-driven reporter line that expresses the membrane-targeted Tomato protein (mT) in the absence of Cre, and membrane-targeted green fluorescent protein (mG) after *Cre*-mediated excision (Muzumdar et al., 2007). Cells in which the *Tcf4* promoter has never been activated fluoresce red at membranes (mTOM+) and are referred to as *Tcf4* lineage-negative (*Tcf4* Lin-). Once the *Tcf4* promoter becomes activated, the cell permanently fluoresces green at membranes (mGFP+) and is referred to as *Tcf4* lineage-positive (*Tcf4* Lin+). Both *Tcf4*-Cre driver alleles represent knock-in, knock-out alleles, so each reporter line is heterozygous for *Tcf4* (Angus-Hill et al., 2011). The *Tcf4^Cre^*allele drives *Cre* expression in response to normal activation of the *Tcf4* promoter. The inclusion of the Neo cassette in the *Tcf4^CreNeo^*line leads to reduced expression from the *Tcf4* locus (Frank et al., 2002), and as a result fewer cells (<1% of the total Tcf4-expressing cells) express *Cre* and become mGFP+ (Angus-Hill et al., 2011). In combination with *Rosa^mTmG^*, *Tcf4^CreNeo^*permanently labeled a subset of *Tcf4*-expressing cells within the colon, including a small number of crypts, and epithelial cells located at the inter-cryptal region in the surface epithelium (Angus-Hill et al., 2011). Previously, we found that homozygous loss of *Tcf4* (generated with two alleles of *Tcf4^CreNeo^*and referred to as *Tcf4^NeoNull^*) caused aberrant proliferation of *Tcf4* Lin+ epithelial cells in the colon crypts of adult animals (Angus-Hill et al., 2011). It appeared that *Tcf4* is normally required to restrict the proliferation of *Tcf4* Lin+ cells in a cell-autonomous, Wnt-dependent manner. However, these hyperproliferative *Tcf4* Lin+ crypts from *Tcf4^NeoNull^*mice, also contained *Tcf4* Lin- cells (Fig. 1B). Labeling with the thymidine analog 5-ethynyl-2′-deoxyuridine (EdU), showed a direct correlation between EdU+ cells and the *Tcf4* Lin- (mTOM+) population (Fig. 1B, yellow arrows). This suggested that loss of *Tcf4* caused hyperplasia of *Tcf4* Lin- cells through a non-cell-autonomous, initially Wnt-independent mechanism. Interestingly, a subset of *Tcf4* Lin- cells were found to be mTOM+, EdU- and mGFP+, suggesting that these cells were transitioning from *Tcf4* Lin- to *Tcf4* Lin+ cells (Fig. 1B, turquoise arrows). These results suggested the possibility of a *Tcf4* Lin- stem-cell population.

**Fig. 1:**
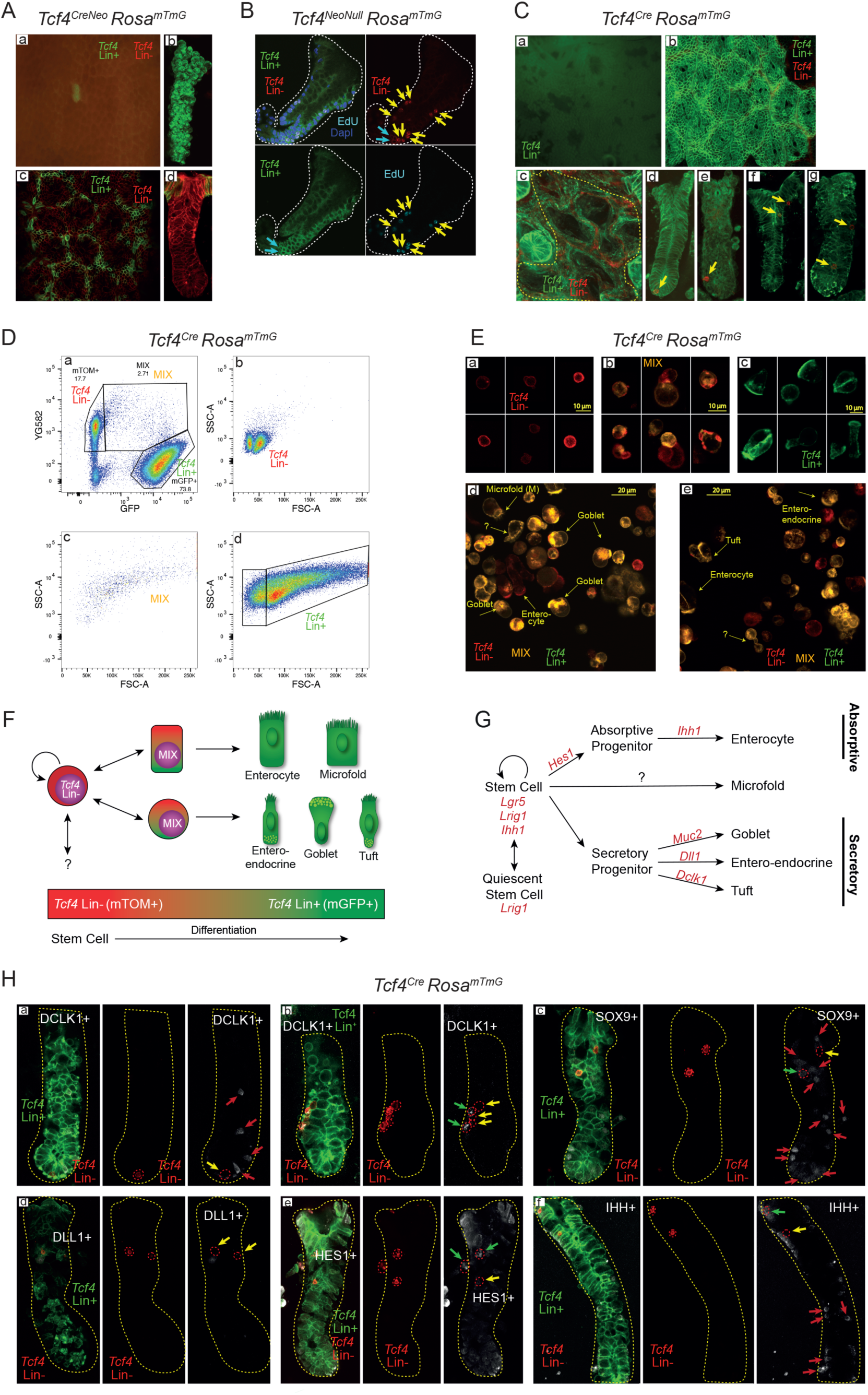
*In vivo* lineage tracing and characterization of *Tcf4* Lin- cells. (A) mGFP marks *Tcf4^CreNeo^*lineage in (Aa,b) full crypts and (Ac,d) differentiated surface epithelium. (B) EdU labeled cells in a *Tcf4^Null^* crypt (mGFP+) isolated from *Tcf4^CreNeo/Lox^* mice are *Tcf4* Lin- (mTOM+) cells (yellow arrows). (C) *Tcf4^Cre^* lineage in surface epithelium (Ca,b) and in isolated crypts (Cc-g). Yellow dotted line in (c) represents a region where crypts have been pushed out of the crypt bed. Yellow arrows mark rare *Tcf4* Lin- cells that appear at different locations throughout the crypt. (D) Isolation of single crypt cells followed by and FACS analysis (Da) demonstrates the presence of (Db) homogeneous Tcf4 Lin- and heterogeneous (Dc) MIX and (Dd) Tcf4 Lin+ cell populations. (E) Tcf4 Lin- descendants have morphological characteristics of secretory and absorptive cell types (scale bar, (Ea-c) 10 μm: (Ed-e) 20 μm). (F) Model depicting *Tcf4* Lineage tracing as stem cells become committed towards absorptive and secretory lineages and (G) the published markers associated with lineage commitment. (H) Co-staining of *Tcf4* Lin+ and Lin- cells and commitment markers. Dotted red lines encircle *Tcf4* Lin- cells. Green arrows denote *Tcf4* Lin- cells that express the commitment marker. Yellow arrows denote *Tcf4* Lin- cells that lack commitment marker expression. Red arrows denote *Tcf4* Lin+ cells that express the commitment marker.

While heterozygous loss of Tcf4 results in no visible phenotype, heterozygous animals exhibit raised neutrophil levels and when combined with Apc mutation develop colon tumors, suggesting defects in epithelial barrier function (Fig. S1; Bice et al., 2017). To further analyze the *Tcf4* Lin- and *Tcf4* Lin+ populations in *Tcf4* heterozygous colon crypts, we utilized the *Tcf4^Cre^ Rosa^mTmG^* line, where *Tcf4* Lin+ cells are found throughout the colon crypt and surface epithelium (Angus-Hill et al., 2011). Colon crypts were isolated using a procedure that reduces contamination from the underlying stromal cells, which are *Tcf4* Lin- (Fig. 1Ca-c, (Nik and Carlsson, 2013), Materials and Methods). Interestingly, *Tcf4* Lin- cells resided at the location commonly occupied by stem cells at the base of the crypt (Fig. 1Cd-e, yellow arrows) and also at locations further up the crypt (Fig. 1Cf-g, yellow arrows).

To further define the *Tcf4* Lin- cell population within the crypt, we dissociated pressure-isolated crypts to obtain a single epithelial-cell suspension (Nik and Carlsson, 2013). Fluorescence-activated cell sorting (FACS) analysis of single cells revealed that 74% of cells were *Tcf4* Lin*+* and 18% were *Tcf4* Lin- (Fig. 1Da). The *Tcf4* Lin- population was comprised of two populations of small cells (Fig. 1Db). In contrast, the *Tcf4* Lin+ population was quite heterogeneous (Fig. 1Dd), as expected from a population comprised of stem cells, transit-amplifying (TA) cells, and secretory and absorptive cells. Interestingly, the analysis revealed a third small population of mTOM+/mGFP+ (“MIX”) cells (Fig. 1Da,c). While the transcription of *mTOM* and *mGFP* genes is mutually exclusive in the *Rosa^mTmG^* reporter mouse, mTOM protein expression lasts for up to 9 days following *Tcf4*-driven CRE expression and subsequent recombination (Muzumdar et al., 2007). Thus, the MIX population represents “newly minted” *Tcf4* Lin+ cells. We further analyzed the fluorescence-sorted cells by confocal microscopy (Fig. 1E) and found that the *Tcf4* Lin- cells have a high nuclear-to-cytoplasmic ratio with no visible internal complexity (Fig. 1Ea; Fig. S2), a phenotype consistent with an intestinal stem cell (Cheng and Leblond, 1974). By contrast, the MIX and *Tcf4* Lin+ populations were quite heterogeneous (Fig. 1Eb-c) since they were highly enriched for goblet, tuft, microfold, and enteroendocrine cells (EEC), as assessed by morphological analysis (Fig. 1Eb,d-e). This demonstrates that the *Tcf4* Lin- population gives rise to all differentiated epithelial cell types, including rare goblet, tuft and microfold cells, and that differentiation into these diverse cell lineages correlates with activation of the *Tcf4* gene.

To analyze the proportion of cycling cells among the three populations, animals were injected with EdU and sacrificed 2 hours following injection. Colon cells were isolated, and cells with EdU incorporation were labeled (Fig. S2A). We found that 2.2% of the *Tcf4* Lin- cells were EdU positive (Fig. S2Ab,f) while the MIX cells and *Tcf4* Lin+ cells had a much higher percentage of cycling cells (20.9% and 20.4%, respectively, Fig. S2Ac,g,d,h). In addition, the non-cycling *Tcf4* Lin- cells (Fig. S2Ba) were slightly smaller than the cycling *Tcf4* Lin- cells (Fig. S2Bb-c). These results indicate that a subset of *Tcf4* Lin- and MIX cells are cycling, a requirement linked to intestinal stem and transit amplifying cells (reviewed in (Stange and Clevers, 2013)).

Our findings suggest that *Tcf4* Lin- cells represent a rare stem-cell/progenitor population in which the commitment to differentiation into multiple cell lineages is initiated by expression of *Tcf4* (Fig. 1F; Angus-Hill et al., 2011). To test this model, we co-stained *Tcf4* Lin- cells within the crypts with antibodies to detect defined epithelial differentiation markers (Fig. 1G). Doublecortin-like kinase 1 (DCLK1) is a tuft cell marker that has also been linked, controversially, to reserve stem-like states (May et al., 2009) and is located at the base of the crypt. Interestingly, *Tcf4* Lin- cells are both DCLK1- (Fig. 1Ha-b, yellow arrows) and DCLK1+ (Fig. 1Hb, green arrows) suggesting that *Dclk1* may be turned on following division of *Tcf4* Lin- cells. Importantly, a population of *Tcf4* Lin+ DCLK1+ cells is also commonly found in crypts (Fig. 1Ha, red arrows). These are likely derived from the Wnt-dependent, LGR5+ CBC cells, since DCLK1+ cells have been shown to co-express LGR5 (Westphalen et al., 2014).

SRY-box 9 (SOX9) is expressed in stem cells and at varying levels in goblet, paneth, EEC and tuft cells (Formeister et al., 2009; Gerbe et al., 2011; Ramalingam et al., 2012; Van Landeghem et al., 2012). While nuclear localization of SOX9 is commonly found in differentiating cells, overexpression or cytoplasmic sequestration of SOX9 can result in reversal of growth arrest in breast cancer cells (Chakravarty et al., 2011). The role of cytoplasmic sequestration of Sox9 in cell growth of colonic or small intestinal cells is currently unknown. Colon crypts isolated from *Tcf4^Cre^ Rosa^mTmG^* animals possess multiple SOX9+ cells throughout the crypt (Fig. 1Hc). Some *Tcf4* Lin- cells are SOX9- (Fig. 1Hc, yellow arrow) and some express cytoplasmic SOX9 (Fig. 1Hc, green arrow). Additionally, cytoplasmic SOX9 is often found in *Tcf4 Lin+* cells located at either the very base of the crypt or in cells closer to the lumen (Fig. 1Hc, red arrows), demonstrating similar SOX9 localization and expression levels in the LGR5+ CBC and in *Tcf4* Lin- cells (Fig. 1Hc, compare red arrows to green arrow). However, nuclear localization of SOX9 is restricted to *Tcf4* Lin+ cells at the crypt base (Fig. 1Hc, red arrows). The combination of *Tcf4* lineage analysis and cytoplasmic localization of SOX9 effectively marks CBC and *Tcf4* Lin- stem cells while nuclear SOX9 marks more differentiated cells.

Delta-like canonical Notch ligand 1 (DLL1) is expressed in the committed, short-lived, TA secretory progenitor population (Fig. 1G) and is able to drive the conversion committed cells to a progenitor population following intestinal damage (van Es et al., 2012). DLL1+ staining was found in crypts among the *Tcf4* Lin+ but not the *Tcf4* Lin- population (Fig. 1Hd). Interestingly, *Tcf4* Lin+ DLL1+ cells were found near *Tcf4* Lin- DLL- cells, raising the possibility that *Tcf4* Lin+ DLL1+ cells arise from the *Tcf4* Lin- population (Fig. 1Hd).

Hes family bHLH transcription factor 1 (HES1) is a downstream target of Notch signaling that is important for both driving absorptive-cell-fate determination (Fig. 1G) and preventing the secretory fate through transcriptional repression (Jensen et al., 2000, Yang, 2001). We found multiple *Tcf4* Lin- and *Tcf4* Lin+ cells throughout the crypt that are HES1+ (Fig. 1He; Fig. S3) validating that HES1 is expressed in *Tcf4* Lin+ cells likely derived from the CBC and in *Tcf4* Lin- cells where it may similarly have a role in cell fate decisions.

Indian hedgehog (IHH) is important for small intestinal stem-cell renewal (Kosinski et al., 2010) and possibly for the differentiation of epithelial cells (van den Brink et al., 2004, Fig. 1G). We therefore examined whole crypts to define whether IHH was expressed in *Tcf4* Lin- cells. We found *Tcf4* Lin- cells that are both IHH+ (Fig. 1Hf, green arrow) and IHH- (Fig. 1Hf, yellow arrow). Additionally, there are multiple *Tcf4* Lin+ cells that are IHH+ (Fig. 1Hf, red arrows). These results suggest a possible role for IHH in *Tcf4* Lin- cell renewal and differentiation.

Taken together, our lineage analysis combined with immunostaining results demonstrates that several known markers of stem cell maintenance and cell fate commitment are expressed in the Tcf4 Lin- cell population. This suggests that this population is a distinct non-CBC stem/progenitor population.

### The *Tcf4* Lin- population is comprised of both stem-cell-like epithelial progenitors and immune-cell populations

Many intestinal stem-cell markers have been defined, including *Lgr5* and *Lrig1* (Barker et al., 2007; Powell et al., 2012; Wong et al., 2012), so we questioned whether they are expressed in *Tcf4* Lin- cells. We isolated single crypt cells from *Tcf4^Cre^ Rosa^mTmG^* animals and used the Drop-seq method (Macosko et al., 2015) for transcriptional profiling of individual FACS-sorted cells from the *Tcf4* Lin- and the MIX population. The cells clustered into 4 distinct populations that are commonly found in the gut based on expression of known markers (Fig. 2A-B; Tab. S1; Tab. S2). The majority (82%) of *Tcf4* Lin- cells were characterized by the presence of immune-cell markers and clustered into one small distinctive group mainly expressing known B cell markers (Fig. 2A-B; Fig. S4A-D) and two larger closely related groups with T-cell characteristics (Fig. 2A-B; Fig. S5). These T-cell clusters were highly similar to intraepithelial lymphocytes (IELs) that are found interspersed between epithelial cells and constitute the front line of defense against pathogens while also maintaining tolerance toward commensal bacteria (Kunisawa et al., 2007). They were identified as natural IELs (nIELs) and induced IELs (iIELs) (Fig. 2A-B; Fig. S4A-D; Fig. S5; Cheroutre et al., 2011; Sheridan and Lefrancois, 2010; Smith et al., 2013). We excluded the possibility of lymphocyte contamination from the stroma since the isolation method specifically selected for the epithelial layer of the large intestine (Nik and Carlsson, 2013). The remaining *Tcf4* Lin- cells (18%) constituted the fourth cluster and were epithelial in origin due to expression of the epithelial markers *Epcam*, *Krt8* and *Krt19* (Fig. 2A, Fig. S4G-H). Because these cells are epithelial, they will from here be designated as “Epi *Tcf4* Lin-“ cells to differentiate them from the immune “IEL *Tcf4* Lin-“ cells. Heat maps further delineated the expression profile differences between the *Tcf4* Lin-populations (Fig. 2B), and additional markers that were unique to the epithelial cluster included the epithelial marker polymeric immunoglobulin receptor gene (*Pigr*), and the oncogene induced transcript 1 gene (*Oit1*) (Fig. 2A-B; Fig. S4E-F). However, further examination by Principal Component Analysis (PCA) revealed heterogeneity within the Epi *Tcf4* Lin- population (Fig. S6A-B), so we analyzed them separately using Seurat (http://satijalab.org/seurat/) and identified three unique Epi *Tcf4* Lin- subclusters (Fig. 2C-D; Tab. S2). The first subcluster was designated “stem cells” due to expression of genes commonly associated with stem cells, proliferation, and development, including *Sox4*, *Anp32b*, *Xist*, *Nop56*, *Nop58* (Fig. 2C-D; Tab. S3). The second subcluster was designated “secretory precursors” due to the expression of secretory lineage markers including *Muc2*, *Spink4*, *Reg4*, *Clca1*, *Fcgbp*, *Agr2*, and *Ang4* (Fig. 2C-D). The third subcluster was designated “absorptive precursors” due to expression of several enterocyte markers including *Gsdmc4*, *Lgals3*, *Aqp4*, *Krt8*, *Krt19*, *Muc3*, and *Car2* (Fig. 2C-D; Tab. S3). Indeed, several absorptive lineage markers were highly expressed in this third population (e.g. fatty acid binding protein 2, *Fabp2*) (Gajda and Storch, 2015) while commonly used markers of differentiated colonocyte/absorptive markers were either not expressed (plasma alkaline phosphatase 1, *Alp1*, data not shown) or only expressed in a few cells (*Ihh*, Fig. 2E). To further analyze these Epi *Tcf4* Lin- subclusters for evidence of lineage commitment to the absorptive or secretory pathways, we examined the expression of *Atoh1*, which marks cells that are committed to the secretory lineage, and *Hes1*, which marks cells that are committed to the absorptive lineage (Toth et al., 2017). Interestingly, we found that *Atoh1* is highly expressed in both the “stem cell” and “secretory precursor” subclusters while *Hes1* is highly expressed in both the “stem cell” and “absorptive precursor” subclusters (Fig. 2F). This indicates that cell-fate decisions likely take place within the Epi *Tcf4* Lin- “stem-cell” subcluster and suggests that the “secretory precursor” and “absorptive precursor” cells are derived from this “stem cell” population.

**Fig. 2:**
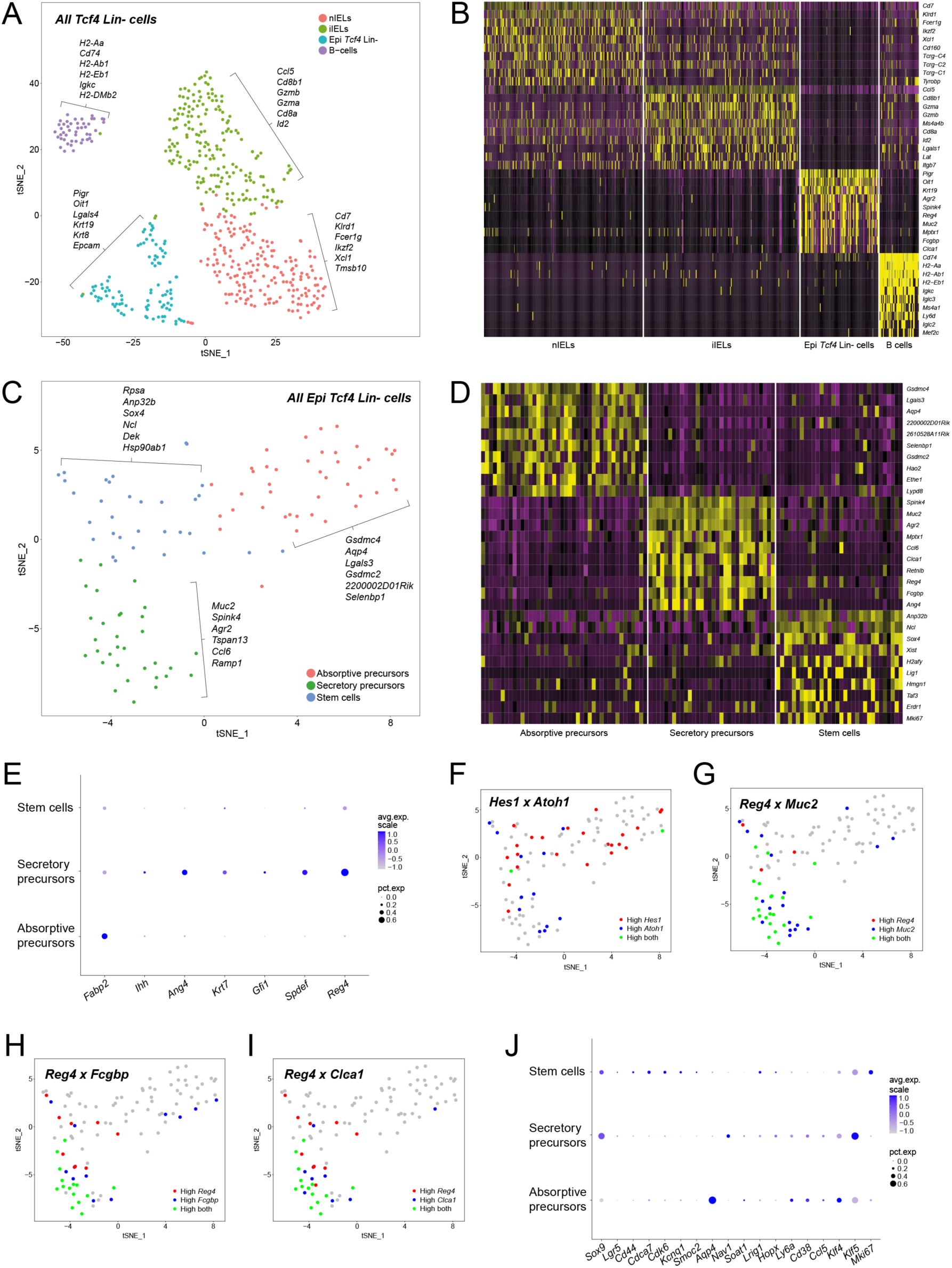
Single cell analysis reveals the presence of three distinctive clusters of stem and precursor cells within Epi *Tcf4* Lin- cells. Analysis of single cell RNA-sequencing data utilizing the R package Seurat. (A) tSNE plot clustering *Tcf4* Lin- cells into four distinctive groups. (B) Top 10 differentially expressed genes of four clusters of *Tcf4* Lin- cells displayed in a heatmap. (C) tSNE plot clustering Epi *Tcf4* Lin- cells into three distinctive groups. (D) Top 10 differentially expressed genes of three clusters of Epi *Tcf4* Lin- cells displayed in a heatmap. (E) Visualization of several absorptive and secretory precursor and differentiation markers via DotPlot, demonstrating average expression level and the percentage of cells expressing the respective gene in each cluster. (F-I) Feature plots displaying the co-expression of two genes simultaneously on the tSNE plot. (F) Co-expression of *Hes1* and *Atoh1*. (G) Co-expression of *Reg4* and *Muc2*. (H) Co-expression of *Reg4* and *Fcgbp*. (I) Co-expression of *Reg4* and *Clca1*. (J) Visualization of several published stem cell markers via DotPlot, demonstrating average expression level and the percentage of cells expressing the respective gene in each cluster.

Intestinal secretory progenitors are known to give rise to goblet, EEC and tuft cells (Fig. 1G; Barker, 2014), and the MIX cell population containing “newly-minted” *Tcf4* Lin+ cells showed evidence of all of these cell types (Fig. 1D). To determine whether the Epi *Tcf4* Lin- secretory precursor population contains cells that are committed to differentiation into these lineages, we further analyzed the expression of secretory-precursor, goblet, EEC and tuft-cell marker genes in the three subclusters. Indeed, we observed expression of two previously described markers of secretory precursors, *Ang4* and *Krt7* (Grun et al., 2015), uniquely in the “secretory precursor” population as well as the goblet-cell differentiation markers *Gfi1* and *Spdef* (Fig. 2I; Noah et al., 2011). Importantly, the EEC marker *Reg4* was expressed by the majority of “secretory precursor” cells (Fig. 2E), but all other published markers of differentiated EECs were not expressed (*Neurog3, Gcg, Pyy, Cck, Sct;* Noah et al., 2011; Nagatake et al., 2014). The *Reg4*+ cells could be further subdivided into those that either co-express, or do not co-express, the goblet-cell markers *Muc2*, *Fcgbp*, and *Clca1* (Fig. 2G-I), suggesting that *Reg4* marks a secretory precursor that eventually gives rise to either EEC or goblet-cell lineages. Finally, *Dclk1* is a marker of tuft cells (Gagliardi et al., 2012) but is not expressed in any of the Epi *Tcf4* Lin- subclusters (Tab. S5). These findings suggest that the Epi *Tcf4* Lin- “secretory precursor” population contains cells that are committed to become goblet cells, but not yet EEC or tuft cells, and that commitment to the latter two lineages may be dependent on the expression of *Tcf4*.

We next questioned the extent to which the Epi *Tcf4* Lin- “stem cell” population differed from other known intestinal stem-cell populations, such as Wnt-dependent *Lgr5*+ CBCs (Barker et al., 2007), *Lrig1*+ stem/progenitor cells (Powell et al., 2012), Sox9^high^ stem cells (Van Landeghem et al., 2012) and Krt19+ stem cells (Asfaha et al., 2015). *Lgr5*+ CBCs are commonly defined by *Lgr5*, *Ascl2*, *Cd44*, *Msi1*, *Olfm4*, and *Sox9* expression (Munoz et al., 2012). We therefore analyzed the single-cell gene sets and found that moderate levels of *Sox9* are frequently expressed primarily in the “stem cell” and “secretory precursor” populations, *Lgr5* and *Cd44* are rarely and minimally expressed, and *Ascl2*, *Msi1*, and *Olfm4* are not expressed (Fig. 2J and data not shown). Additional markers of *Lgr5*+ CBC cells, including *Cdca7*, *Cdk6*, *Kcnq1* and *Smoc2*, were modestly expressed in the Epi *Tcf4* Lin- “stem cell” population, while expression levels of *Aqp4, Nav1*, and *Soat1* varied and were not restricted to these cells (Fig. 2J; (Munoz et al., 2012)). Notably, many markers specific for the Epi *Tcf4* Lin- “stem cell” population (Fig. 2C) have not been described as markers of *Lgr5*+ CBCs, so we examined expression of the quiescent-stem-cell markers *Bmi1*, *Tert*, *Hopx*, and *Lrig1* (reviewed in (Smith et al., 2016)). Of these markers, only *Lrig1* and *Hopx* were expressed, and only very rarely within the Epi *Tcf4* Lin- “stem cell” population (Fig. 2J). However, several characterized Lrig1+ cell markers, including *Ly6a*, *Cd38*, *Ccl5*, and others are expressed in subsets of Epi *Tcf4* Lin- cells. Sox9^high^ stem cells are enriched for quiescent stem-cell markers *Bmi1*, *Hopx*, *Dclk1*, and for (EEC) markers (Van Landeghem et al., 2012). While some Epi *Tcf4* Lin- cells express low levels of *Sox9*, very few cells express *Hopx*. Interestingly, approximately 80% of “absorptive precursors” expressed *Krt19* at high levels. Sixty percent of the “stem cell” cluster and 40% of “secretory precursors” expressed lower levels of *Krt19*, perhaps only coincident with expression of *Krt19* in 3% of colon cells (Asfaha et al., 2015). These findings indicate that the Epi *Tcf4* Lin-“stem cell” population can be identified by a unique gene signature with limited overlap of expression of known intestinal-stem-cell markers.

### The *Tcf4* Lin- “stem cell” population gives rise to *Tcf4* Lin- “secretory precursor” and “absorptive precursor” cell lineages

To further analyze whether the Epi *Tcf4* Lin- “stem cell” population is likely to give rise to the associated “absorptive precursor” and “secretory precursor” populations, we performed differential expression and time-series analysis on single crypt cells using Monocle (http://cole-trapnell-lab.github.io/monocle-release/). This method allows for the analysis of cells progressing in pseudotime through the process of cell fate determination or differentiation. Subcluster-specific markers *Sox4* (“stem cell”), *Muc2* (“secretory precursors”) and *Lgals3* (“absorptive precursors”) that were identified in Fig. 2C were plotted and analyzed using Monocle, and the existence of the three distinctive cell states was consistent with Seurat analysis (Fig. 3A-D, Fig. S7). *Sox4*+ cells marking the “stem cell” population were identified at the earliest point in time before the branch point indicating that cell-fate decisions are made within this group to either follow the *Muc2*+ secretory differentiation pathway or the *Lgals3*+ absorptive differentiation pathway (Fig. 3A-D). Interestingly, cells with detectable *Tcf4* and *Apc* transcripts were first observed predominantly past the branch point, consistent with roles for both genes in cell commitment and differentiation (Andreu et al., 2005; Heuberger et al., 2014; Fig. 3E-F). These findings are consistent with a model whereby the *Sox4*+ *Pigr*+ *Tcf4* Lin- cells represent a stem-cell population that gives rise to “secretory precursor” and “absorptive precursor” Epi *Tcf4* Lin- cells.

**Fig. 3:**
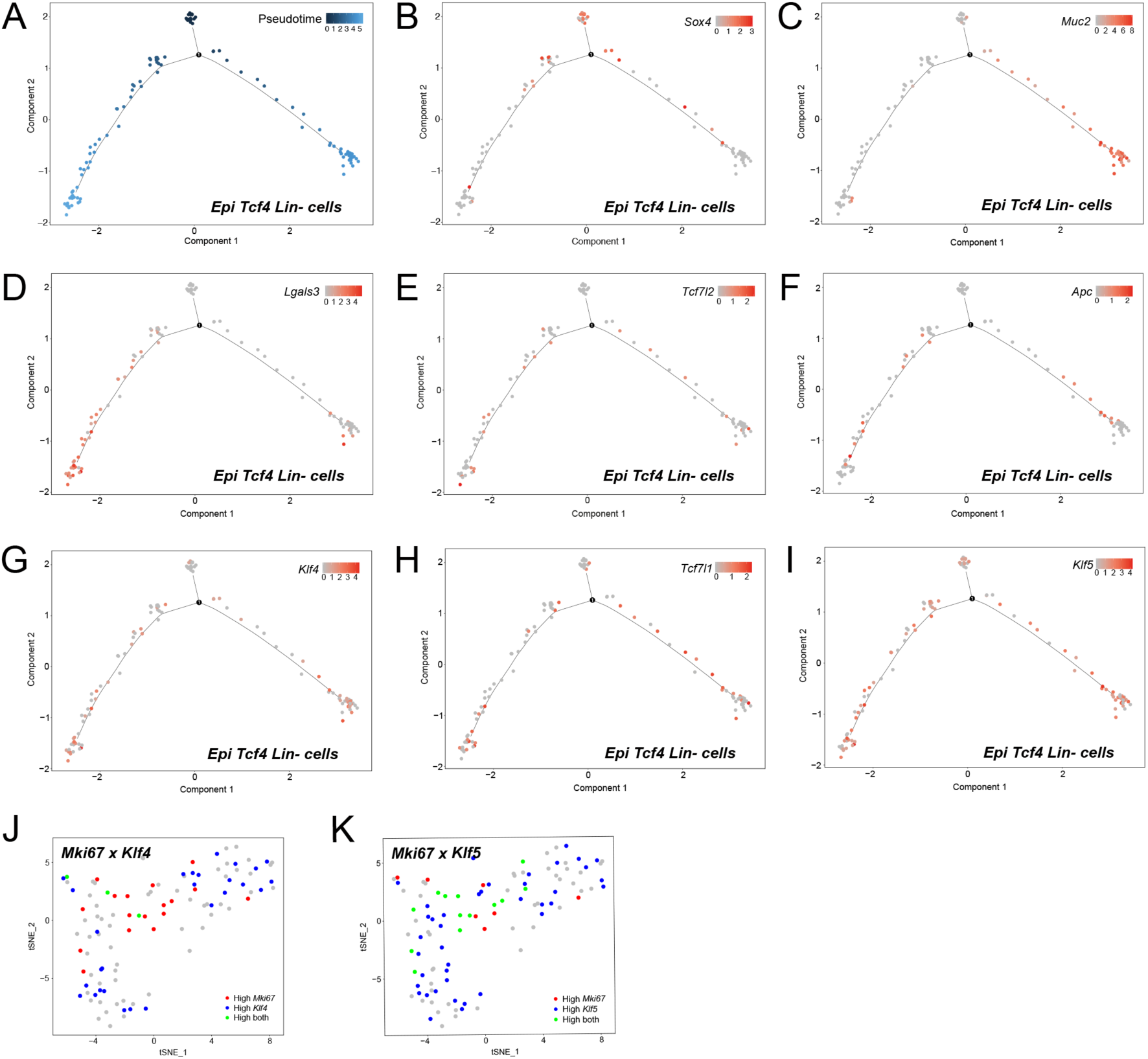
Organization of Epi *Tcf4* Lin- cells in pseudotime confirms that a stem cell cluster gives to either an absorptive or a secretory cell fate. Analysis of the pseudotime properties of Epi *Tcf4* Lin- cells by employing the monocle algorithm. (A) Single cell trajectory analysis, displaying pseudotime, places each cell along the trajectory depending on the amount of transcriptional change each cell has undergone. (B-I) Single cell trajectories displaying the expression level of a marker and placing cells in pseudotime. (B) *Sox4*, a distinctive marker of the Stem cell cluster. (C) *Muc2*, a distinctive marker of Secretory precursors. (D) *Lgals3*, a distinctive marker of Absorptive precursors. (E) *Tcf7l2*. (F) *Apc*. (G) *Klf4*. (H) *Tcf7l1*. (I) *Klf5*. (J-K) Seurat analysis: Feature plots displaying the co-expression of two genes simultaneously on the tSNE plot. (J) Co-expression of *Mki67* and *Klf4*. (K) Co-expression of *Mki67* and *Klf5*.

The Krüppel-like factors (KLF) *Klf4* and *Klf5* are highly expressed in the gastrointestinal tract (McConnell and Yang, 2010). *Klf5*, a transcription factor involved in self-renewal of embryonic stem cells, is expressed in actively proliferating cells of the intestinal crypt, including both *Lgr5*+ CBCs and TA cells (Nandan et al., 2015). In contrast, *Klf4*, an inhibitor of cell cycle progression, is highly expressed in cells that are differentiating toward the secretory and absorptive cell lineages (McConnell and Yang, 2010). Since low levels of *Klf4* mark the proliferative compartment, its overall expression pattern is similar to *Tcf4* (Angus-Hill et al., 2011; Flandez et al., 2008; McConnell and Yang, 2010).

*Tcf4* (*Tcf7l2*) and *Klf4* showed similar expression patterns (Fig. 3E,G), marked by a later onset of gene expression as the cells progressed through pseudotime, while the expression pattern for *Tcf3* (*Tcf7l1*) resembled the pattern for *Klf5* (Fig. 3H,I), in that genes were expressed earlier on in pseudotime. Moreover, *Klf5*, but not *Klf4*, in Epi *Tcf4* Lin- cells overlap with the proliferative marker *Mki67* expression (Ki67, Fig. 3J,K). Expression of *Sox4*, *Mki67*, and *Klf5* marks the “stem cell” cluster. As *Sox4* expression is shut off, *Klf4* and *Tcf4* are turned on in both secretory and absorptive precursors and together act as markers of committed cells. The *Sox4*+ Epi *Tcf4* Lin- stem cell therefore appears to be poised to switch from proliferation to lineage commitment and differentiation.

### The epithelial *Tcf4* Lin- cell population gives rise to cells that are important for colon crypt barrier repair

Single-cell sequencing revealed that *Tcf4* Lin- cells represent a heterogeneous population of Epi *Tcf4* Lin-progenitors, IELs and B cells (Fig. 2A,B). We set out to remove non-epithelium-derived cells from the *Tcf4* Lin- population, in order to analyze the Epi *Tcf4* Lin- cells separately from the other IEL populations. The *Cd2^iCre^* line is designed to drive conditional recombination of Cre recombinase (iCre; an optimized variant) in all T-cell and B-cell progenitors within the colon-crypt lymphocyte population (de Boer et al., 2003). To first analyze the *Cd2* Lin+ (IEL, mGFP+) population in the colon crypt, we combined the *Cd2^iCre^* line with *Rosa^mTmG^*, and found Cd2+ mGFP+ cells in all crypt locations: at the base, +4 position, middle, and top (Fig. S8Aa-d). Importantly, the crypt isolation method (see Fig. 1C) leaves many lymphocytes within the stroma (Fig. S8Ae). We found that 68% of isolated *Cd2^iCre^ Rosa^mTmG^* crypts contained *Cd2* Lin+ cells, with greater than 50% of these cells located in the mid- to upper-crypt region (Fig. S8Ba-b; Tab. S4). FACS analysis of single cells from isolated *Cd2^iCre^ Rosa^mTmG^* crypts revealed that *Cd2* Lin+ mGFP+ cells comprised about 1% of total crypt cells (Fig. S8Ca), and over 50% of these cells were positive for the T-cell marker *CD8a* (Fig. S8Cb). These findings further support the conclusion that IELs represent one of the *Tcf4* Lin- cell populations in the colon crypts, and importantly, the *Cd2^iCre^ Rosa^mTmG^* mouse line allows for the exclusion of these cells from subsequent analyses.

To characterize the epithelial *Tcf4* Lin- cells in isolation from the IEL *Tcf4* Lin- populations, we generated *Cd2^iCre^ Tcf4^Cre^ Rosa^mTmG^* animals, where cre expression from *Cd2^iCre^*and *Tcf4^Cre^* loci drive cre-mediated excision of mTOM in either *Cd2* Lin+ and/or *Tcf4* Lin+ cells. Importantly, expression of mGFP in *Cd2* Lin+ *Tcf4* Lin- lymphocytes enables cell separation from mTOM expressing Epi *Tcf4* Lin- cells. Tcf4 Lin- cells of epithelial origin therefore remain mTOM+ due to the absence of *Cd2^iCre^* expression in this epithelial cell type. Similar to the *Cd2* Lin+ population (Fig. S8Aa-d), Epi *Tcf4* Lin- cells were present at all locations within the crypt: the crypt base, +4 position, the middle, and the top (Fig. 4Aa-f). Interestingly, rare crypts contained a large number of Epi *Tcf4* Lin- cells, as well as MIX cells, and were often shortened and contained fewer cells (Fig. 4Aa and 4Ba), suggesting that Epi *Tcf4* Lin- cells may help to expand damaged crypts. Excluding these rare crypts, an average of 14% of isolated crypts contain an Epi *Tcf4* Lin^-^ cell with similar numbers found in the mid- to upper-crypt region and fewer at the +4 position and crypt base (Fig. 4Bb-c; Tab. S4).

Forty percent of the Epi *Tcf4* Lin- “stem cell” subcluster expressed Ki67 (*Mki67*, Fig. 2J), suggesting that many of these cells are proliferative (Schluter et al., 1993). To determine whether the Epi *Tcf4* Lin- population is actively engaged in the cell cycle, *Cd2^iCre^ Tcf4^Cre^ Rosa^mTmG^* mice were collected 2 hours following EdU injection, colonic crypts were isolated, and the single-cell population was analyzed by FACS. Calcein violet staining revealed that the majority of isolated cells from mTOM+, MIX, and mGFP+ populations were viable (Fig. 4Ca-d). The Epi *Tcf4* Lin- population comprised fewer than 2% of total colon-crypt cells, while the percentage of MIX cells was 1% (Fig. 4Ce). Next, EdU-labeled cells were analyzed by FACS. Nine percent of Epi *Tcf4* Lin- cells were found to be in S-Phase and 6% in G2/M phase (Fig. 4Cf), indicating a high level of proliferation of this population. MIX cells showed 14% in S-phase and 13% in G2/M (Fig. 4Cg), suggesting that this population may consist of progenitor cells similar to TA cells (Barker et al., 2008a). As expected, the mGFP population, a subset of which is comprised of rapidly dividing *Lgr5+* CBC and TA cells, is also proliferative (Fig. 4Ch,f; (Barker et al., 2008a)). Many of the proliferative Epi *Tcf4* Lin- and EdU+ MIX were round and 8-10 µm in size and had a high nuclear to cytoplasmic ratio (Fig. 4D-E), a morphological phenotype similar to stem cells (Cheng and Leblond, 1974). Therefore, these rare Epi *Tcf4* Lin- “stem cells” are actively proliferating and have the morphological characteristics of stem cells.

**Fig. 4:**
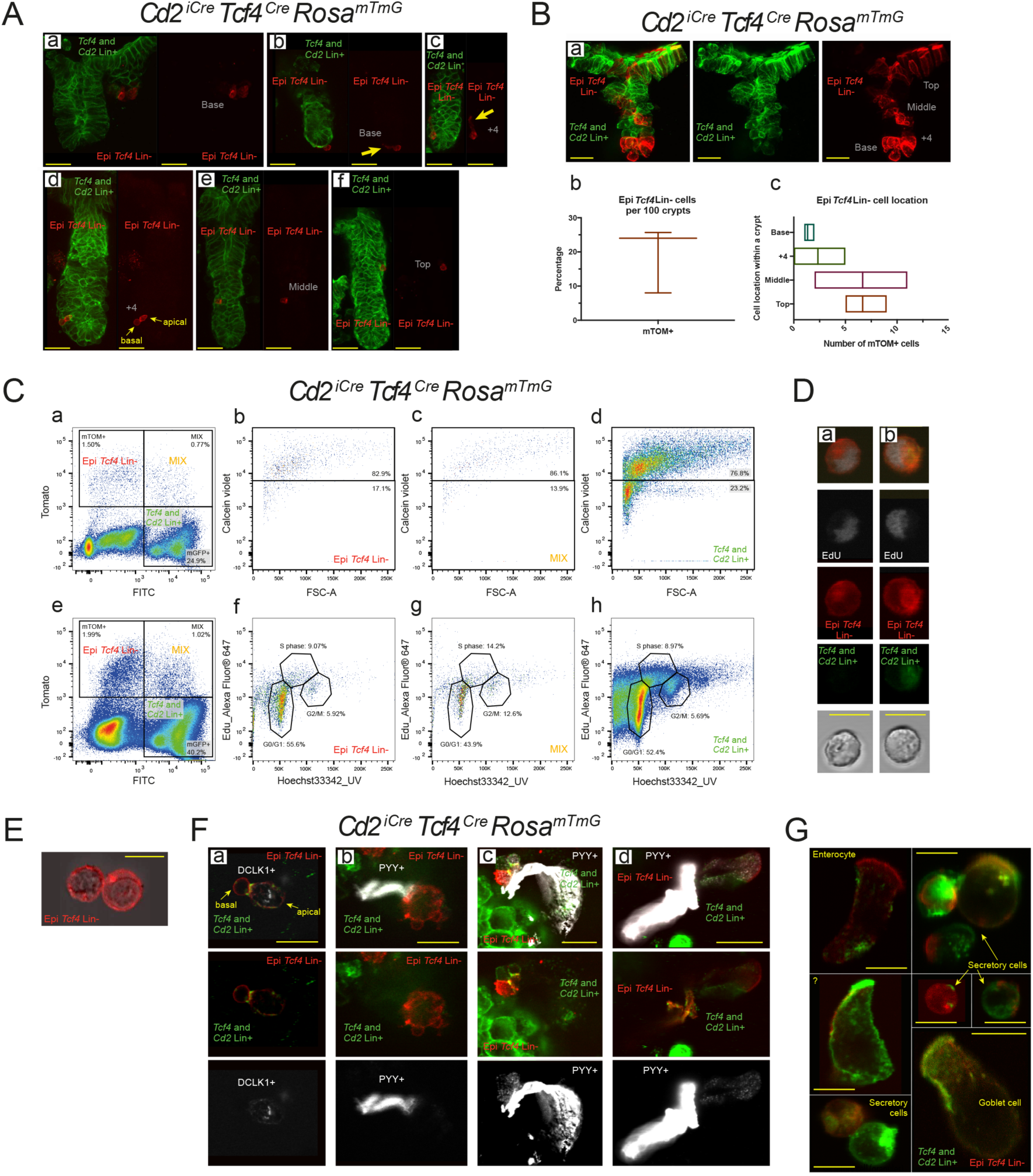
Characterization of the Epi Tcf4 Lin- population revealed a proliferative stem cell population that gives rise to secretory and absorptive lineages. (A-B) Confocal Images, cell number, and crypt location of Epi Tcf4 Lin- and MIX cells (scale bar, 50 μm). Yellow arrows mark unique cellular characteristics (C) EdU labeling and FACs analysis for cell viability (Ca-d) and proliferation (Ce-h) of Epi *Tcf4* Lin-, MIX and *Tcf4* and *Cd2* Lin+ cell populations. (D-E) A representative example of morphological analysis of fluorescence of Epi *Tcf4* Lin- and MIX cell populations combined with (D) fluorescent EdU+ and (E) differential interference contrast (Nomarski DIC; scale bar, 10 μm). (F-G) Characterization of Epi *Tcf4* Lin- and MIX populations (F) by co-staining with antibodies to known cell markers DCLK1 (Tuft) and PYY (Enteroendocrine L-cells) or (G) by morphology (scale bar in (F) is 20 μm and in (G) is 10 μm). Note that the MIX population is where mGFP and mTOM are labeling membranes in the same cell but are not overlapping.

The finding that the Epi *Tcf4* Lin- cell population is rare and appears more frequently in shortened crypts (Fig. 4Aa and 4Ba), suggests that this population may be important for barrier repair following injury. Several epithelial cell types are specifically activated following gut barrier damage, including the tuft cell and the enteroendocrine L-cell (Lebrun et al., 2017; Westphalen et al., 2014). To define whether tuft or L-cells can be derived from Epi *Tcf4* Lin- cells, we stained isolated crypts and single cells with either tuft (DCLK1) or L-cell (PYY) markers. Tuft cells have a bulge at the base and a narrow apical portion with microvilli that project into the crypt lumen (Bjerknes et al., 2012; McKinley et al., 2017; Sato, 2007). We found that DCLK1 is expressed in MIX cells that have the morphological features of tuft cells (Fig. 4Ad, 4Fa, (McKinley et al., 2017)). In contrast, *Dclk1* expression was not found in Epi *Tcf4* Lin- cells (Tab. S5). In the colon, L-cells are spindle-shaped cells that secrete PYY and intercalate between epithelial cells (Bohorquez et al., 2011). A subset of colonic Epi *Tcf4* Lin- derived cells had basal processes that extended across the crypt and remain attached to the epithelium, although they did not morphologically resemble colonic enteroendocrine L-cells (Fig. 4Ab-c, arrows). This phenotype is actually reminiscent of L-cells that reside in the ileum and have basal pseudopod-like processes called neuropods that extend between the mucosal epithelium and the lamina propria (Bohorquez et al., 2014). Based on morphology, therefore, a subset of colonic Epi *Tcf4* Lin- derived cells are similar to ileal enteroendocrine L-cells (Bohorquez et al., 2011). These cells are derived from the Epi *Tcf4* Lin-population as Epi *Tcf4* Lin- cells fail to express PYY or *Glp1*, another L-cell marker (by gene expression analysis; Tab. S5), while Epi *Tcf4* Lin- derived MIX cells contain pseudopod-like processes that are PYY-positive (Fig. 4Fb-d). Finally, several other Epi *Tcf4* Lin- derived (MIX) cell types are apparent in isolated cells from colonic crypts, including goblet cells and other granule-containing secretory cells, colonocytes and other unknown cell types (Fig. 4G). These results suggest that the *Tcf4* Lin- cell population gives rise to cells that are important for colon crypt barrier repair.

### Epithelial *Tcf4* Lin- cells are likely recruited to the wound bed during the wound repair process

We next evaluated whether Epi *Tcf4* Lin- cells are recruited as part of colonic wound repair by utilizing a wounding technique that uses visual endoscope guidance and mucosal biopsy to produce discrete mucosal lesions in the colons of *Cd2^iCre^ Tcf4^Cre^ Rosa^mTmG^* mice (Video S1; Fig. 5A-B; Seno et al., 2009). At 3 days post-injury, post-mitotic wound-associated epithelium (WAE, Fig. 5Ca; turquoise arrows) and a single-cell layer of epithelium derived from crypts adjacent to the injury site (Fig. 5Ca; magenta arrows) initially cover the wound bed, as expected (Seno et al., 2009). Both the WAE and the single-cell layer of epithelium are *Tcf4* Lin+ (Fig. 5Ca), indicating that both populations are derived from cells that had activated the *Tcf4* promoter prior to injury. However, co-staining with an antibody to epithelial cell adhesion molecule (EPCAM), a marker of epithelium, revealed *Tcf4* Lin- epithelial (EPCAM+ mTOM+) and newly-minted *Tcf4* Lin+ epithelial (EPCAM+ MIX) populations in the crypts adjacent to the injury boundary, which appear to be expanding into the wound bed (Fig. 5Cb; yellow arrows). A MIX PYY+ population in the region of the crypts is also expanding into the wound bed (Fig. 5Db; red arrows), which is consistent with a role for PYY-expressing MIX cells in wound repair.

**Fig. 5:**
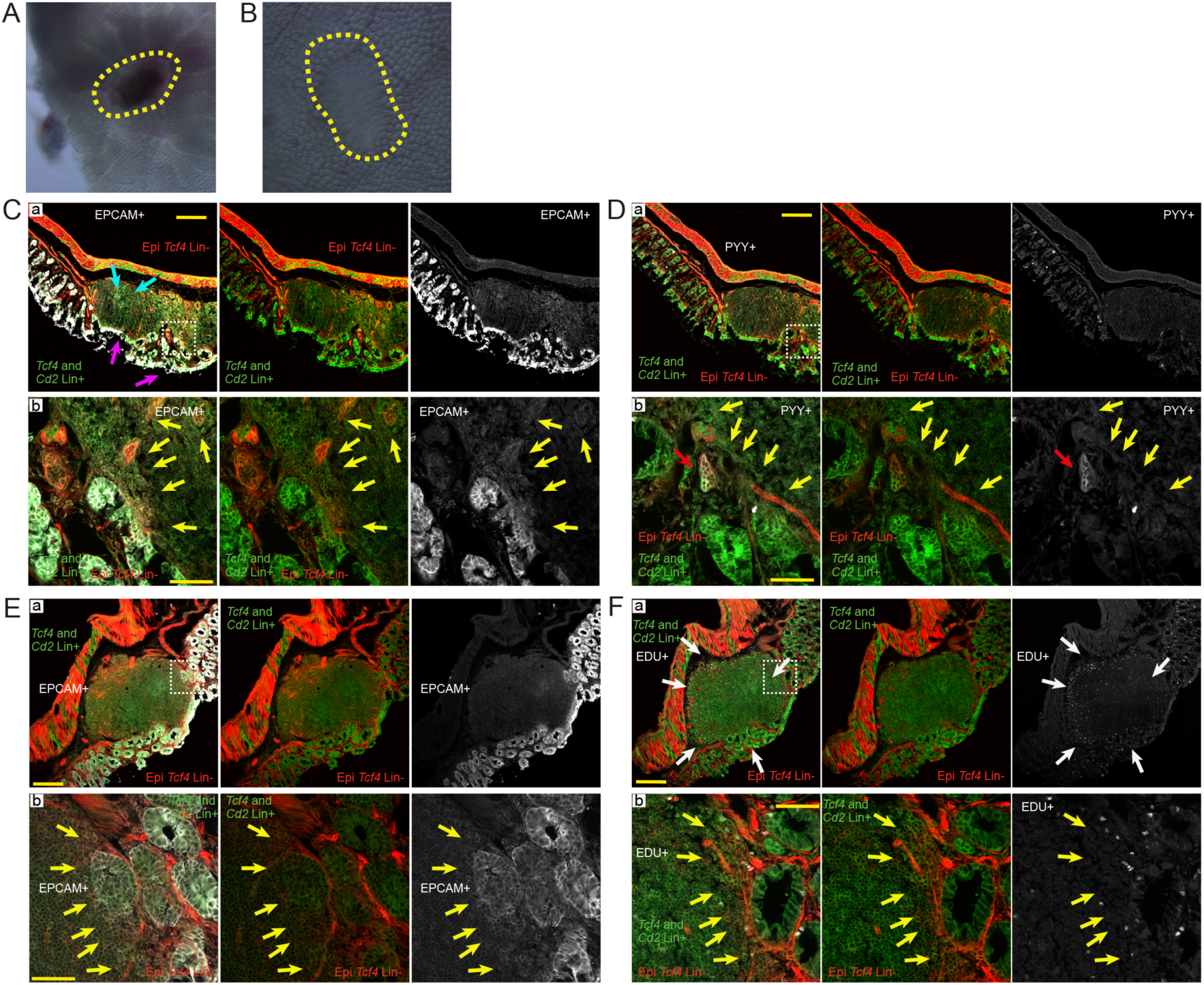
Mechanical damage in colon crypts results in recruitment to the wound bed and proliferation of Epi *Tcf4* Lin- cells for crypt restoration. Analysis of the wound bed at 3 days (A,C-D) and 5 days (B, E-F) post-endoscopic biopsy. (A-B) Whole mount microscopy of wound bed at 3 days (A) and 5 days (B) post-damage. Yellow lines encircle the wound. (C-D) 3 days following damage, adjacent tissue sections were stained with EPCAM and PYY and (E-F) 5 days following damage sections were stained with EPCAM and EdU, and analyzed along with Tcf4 lineage status. White boxes in Ca,Da,Ea,Fa (scale bar is 200 μm) denote regions of magnification in Cb,Db,Eb,Fb (scale bar is 50 μm). Turquoise arrows mark the WAE, and magenta arrows mark adjacent crypts that initially cover the wound bed. Yellow arrows mark the injury boundary. Red arrows mark PYY positive MIX cells and white arrows mark EdU+ regions of proliferation.

Epithelial proliferation has previously been shown to be limited to cells within crypts that are adjacent to the wound bed following colonic damage, and not within the WAE (Seno et al., 2009). To define whether epithelial *Tcf4* Lin- and epithelial MIX cells adjacent to the wound bed are proliferative, we analyzed tissue collected at 5 days post-injury for EPCAM to mark epithelial regions (Fig. 5E) and in adjacent sections for EdU incorporation (Fig. 5F). Similar to previous studies (Seno et al., 2009), EdU incorporation was primarily found in cells adjacent to the wound bed (Fig. 5Fa; White arrows). Importantly, these regions were EPCAM+ *Tcf4* Lin- and EPCAM+ MIX and were situated where crypts are expanding to fill the wound bed (Fig. 5Eb and Fig. 5Fb, yellow arrows). This finding suggests that epithelial *Tcf4* Lin- and newly transitioned *Tcf4* Lin+ cells adjacent to the wound bed promote the expansion of epithelial crypts as part of the wound repair process.

### Colon tumors originate from different cell populations depending on the nature of the *Apc* mutation

It has been proposed that colon tumors arise from proliferative progenitor cells or stem cells within the colon crypt, but whether tumor cells can originate from multiple stem cell pools or even differentiated cell types is unknown (Barker et al., 2009). We therefore set out to test whether mutation of the colon tumor suppressor gene *Apc* could give rise to tumors from both the *Tcf4* Lin- and *Tcf4* Lin+ populations (Fig. 6A-C). To address this question, *Tcf4^CreNeo^*and *Tcf4^Cre^* drivers (Fig. 1A,C) were used to drive conditional deletion of the entire *Apc* gene (*Apc^LoxEx1-15^*, Fig. 6A, Cheung et al., 2010) or truncation of *Apc* through deletion of exon 5 (*Apc^LoxEx5^*, Fig. 6B, Knock-out Mouse Project (KOMP)), specifically within the *Tcf4* lineage. *Hprt^Cre^*-mediated germline deletion or truncation of *Apc*, as well as the *Apc^Min^*(truncation at amino acid 850) mouse line, were also analyzed for comparison. To verify that recombination rates were similar between *Apc^LoxEx1-15^* and *Apc^LoxEx5^* animals, we performed a genotyping analysis of tissue isolated from the colons of recombined animals (Fig. 6D). This is important since the *Apc* locus is relatively large, and the *LoxP* sites in *Apc^LoxEx1-15^* animals are about 200 kb apart, while the *LoxP* sites in the *Apc^LoxEx5^* allele are only a few hundred base pairs apart. Not surprisingly, very little *Apc* is recombined in colons isolated from *Tcf4^CreNeo^* animals, given the hypomorphic nature of this allele (Fig. 6D, Angus-Hill et al., 2011). However, similar levels of *Tcf4^Cre^*-dependent *Apc* recombination are found in *Apc^LoxEx1-15^* and *Apc^LoxEx5^* animals indicating that recombination was efficient despite the large distance between *LoxP* sites in the *Apc^LoxEx1-15^* allele (Fig. 6Da compared to 6Db).

**Fig. 6:**
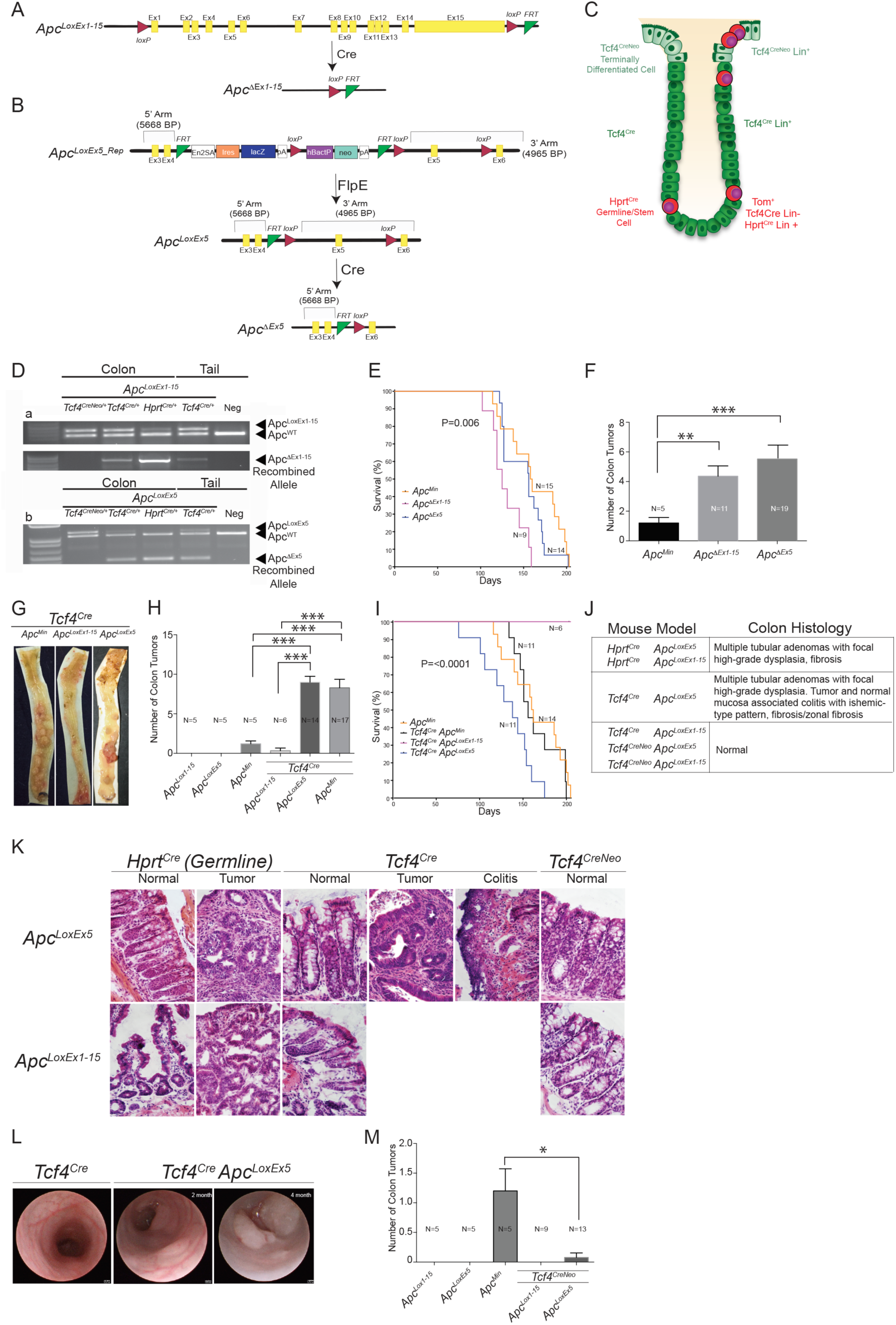
Colon tumors initiate following truncation and not deletion of *Apc* in Epi *Tcf4* Lin+ cells. Structure of the (A) *Apc^LoxEx1-15^* and (B) *Apc^LoxEx5^* alleles. (C) A cartoon depicting the crypt localization patterns of *Tcf4^Cre^* expression, and compared to *Hprt^Cre^* Germline patterns (in all cells), and *Tcf4^CreNeo^* in stem cells of rare crypts and differentiated surface epithelium. (D) Genotyping analysis conducted on colon and tail tissue collected from (Da) *Apc^LoxEx1-15^* and (Db) *Apc^LoxEx5^*and negative control mice following Cre-dependent recombination. (E) Survival Curves for *Apc^Min/+^*, *Apc***^Δ^***^Ex1-15^ (Hprt^Cre^ Apc^LoxEx1-15/+^*) and *Apc***^Δ^***^Ex5^ (Hprt^Cre^ Apc^LoxEx5/+^*) mice. Median survival days were 159 (n=14), 125 (n=9), and 156 (n=14), respectively. P values by chi-square statistic were calculated using the log rank Mantel-Cox test. (G-H) Whole mount colons (G) and colon tumor counts (H) in colons isolated from 4-month-old *Apc^LoxEx5^ Tcf4^Cre^ and Apc^LoxEx1-15^ Tcf4^Cre^* animals. ***p = 0.0005 using two–sample t- test with Welch correction. Error bars were calculated using SEM. (I) Survival Curves for *Tcf4^Cre/+^ Apc^Min/+^*, *Tcf4^Cre/+^ Apc^LoxEx5/+^*and *Tcf4^Cre/+^ Apc^LoxEx1-15^* with *Apc^Min/+^* animals for comparison. Median survival days were 154 (n=11; *Tcf4^Cre/+^ Apc^Min/+^*) and 136 (n=11; *Tcf4^Cre/+^ Apc^LoxEx5/+^*). At the end of the 8-month study, all *Tcf4^Cre/+^ Apc^LoxEx1-15/+^* animals survived and had no visible phenotypes and were censored from the study. P values by chi-square statistic were calculated using the log rank Mantel-Cox test for greater than 3 groups. (J) Histological analysis summary and (K) histology of colon tissue sections of colons isolated from *Apc^LoxEx1-15^*and *Apc^LoxEx5^* conditionally deleted by *Hprt^Cre^ Tcf4^Cre^* and *Tcf4^CreNeo^* driven recombination. (L) Representative endoscopy of the colon in 2 month and 4-month-old *Tcf4^Cre/+^ Apc^LoxEx5/+^* animals. (M) colon tumor counts in colons isolated from 4-month-old *Apc^LoxEx5^Tcf4^CreNeo^ and Apc^LoxEx1-15^ Tcf4^CreNeo^* animals. *p = 0.05 using two–sample t- test with Welch correction. Error bars were calculated using SEM.

Germline heterozygous *Apc* deletion has been shown to be more detrimental in regard to lifespan and colon tumorigenesis than mice harboring a heterozygous *Apc^Min^* mutation (Cheung et al., 2010). To determine whether this effect was due to *Apc* whole-gene deletion versus truncation, *per se*, we compared lifespan and colon tumor load between the three germline *Apc* mutant lines (*Hprt^Cre^ Apc^LoxEx5/+^*, *Hprt^Cre^ Apc^LoxEx1-15/+^*and *Apc^Min/+^*). Similar to previous findings, we found that mice with an *Apc* germline deletion (*Hprt^Cre^ Apc^LoxEx1-15/+^*) exhibited decreased survival (by 28%, Fig. 6E) and a 3-fold higher colon tumor burden (Fig. 6F) than *Apc^Min/+^*animals (Cheung et al., 2010). However, we found that while *Hprt^Cre^ Apc^LoxEx5/+^* and *Apc^Min/+^*mutant animals exhibited similar median survival (Fig. 6E), *Hprt^Cre^ Apc^LoxEx5/+^* animals had 4.3-fold more colon tumors than *Apc^Min/+^* animals (Fig. 6F). Similarly, *Hprt^Cre^ Apc^LoxEx5/+^* animals also developed more small intestinal tumors (SI; Fig. S9B). Thus, germline deletion of *Apc* consistently decreased survival of mice compared to germline truncation of Apc, but did not necessarily drive the formation of more colon tumors.

To determine whether *Apc* deletion and *Apc* truncation can drive colon tumorigenesis from the *Tcf4* Lin+ CBC stem-cell derived population in a similar manner, we compared lifespan and colon tumorigenesis in *Tcf4^Cre^ Apc^LoxEx1-15/+^* and *Tcf4^Cre^ Apc^LoxEx5/+^* animals. As a reminder, both lines are heterozygous for *Tcf4* and lead to disruption of *Apc* after activation of the *Tcf4* promoter. We recapitulated our previous finding (Angus-Hill et al., 2011) that germline *Tcf4* haploinsufficiency exacerbates *Apc^Min^*-mediated colon tumorigenesis (Fig. 6G-H) with minimal effects on overall survival (Fig. 6I). Similarly, truncation of *Apc* specifically in *Tcf4*-expressing cells (*Apc^LoxEx5/+^ Tcf4^Cre/+^*) causes 8-fold more colon tumors and more SI tumors than seen in *Apc^Min/+^* animals (Fig. 6H, Fig. S9C), while median survival is drastically reduced (Fig. 6I). Interestingly, *Apc^LoxEx5/+^ Tcf4^Cre/+^*animals show poorer survival than animals with *Hprt^Cre^* driven germline truncation of *Apc^LoxEx5/+^* (compare blue line in Fig. 6I with blue line in Fig. 6E). Histological examination of colon tissue isolated from both *Hprt^Cre^ Apc^LoxEx5/+^* and *Tcf4^Cre/+^ Apc^LoxEx5/+^* animals demonstrated multiple tubular adenomas with focal high-grade dysplasia but only *Tcf4^Cre/+^ Apc^LoxEx5/+^*mice exhibited colitis with ischemic-type pattern and fibrosis that is localized to both tumor and normal mucosa (Fig. 6J-L). Finally, unlike *Tcf4^Cre/+^ Apc^LoxEx5/+^* animals, *Tcf4^Cre/+^ Apc^LoxEx1-15/+^* mice rarely develop colon (mean 0.3 +/- 0.3) or SI tumors and survive up to twice as long (8 months) without visible phenotypes (Fig. 6H,I; Fig. S9C). This surprising result demonstrates that heterozygous deletion of *Apc* in *Tcf4* Lin*+* cells (*Tcf4^Cre/+^ Apc^LoxEx1-15/+^)* is not sufficient for tumorigenesis and suggests that colon tumors driven by germline deletion of *Apc* (*Hprt^Cre^ Apc^LoxEx1-15/+^*) originate from *Tcf4* Lin-cells. Moreover, since conditional heterozygous deletion of *Apc^LoxEx1-15^* or truncation of *Apc^LoxEx5^* in the *Cd2^iCre^* lineage does not lead to the development of colon tumors (Tab. S6), our data suggest that deletion of *Apc* drives colon tumorigenesis from the Epi *Tcf4* Lin- stem-cell derived population.

It has previously been shown that tumors arise from stem cells located at the base of the crypt (Barker et al., 2009). However, it is unclear whether tumors arise following conditional deletion of *Apc* predominantly in differentiated intercryptal epithelial cells. To address this possibility, we utilized the *Tcf4^CreNeo^* allele, which mainly drives recombination in a subset of *Tcf4*-expressing, differentiated intercryptal epithelial cells (Angus-Hill et al., 2011; Fig. 1). Figure 6M demonstrates that neither deletion (*Tcf4^CreNeo/+^ Apc^LoxEx1-15/+^*) nor truncation (*Tcf4^CreNeo/+^ Apc^LoxEx5/+^*) of *Apc* within these *Tcf4*-expressing differentiated cells is sufficient to drive the rapid development of colon tumors. Interestingly, both *Apc* mouse lines develop small intestinal tumors, however with vastly more duodenal tumors in *Apc^LoxEx5/+^*animals compared to *Apc^LoxEx1-15/+^* with the *Tcf4^CreNeo^*driver (Fig. S9D). These findings demonstrate that disruption of the *Apc* gene within *Tcf4^CreNeo^* expressing epithelial cells, in combination with germline heterozygosity for *Tcf4*, is sufficient for the development of tumors in the small intestine but not the colon.

### Proposed model for epithelial Tcf4 Lin- cell function in wound repair and Apc allele-specific tumor initiation

Our data suggest a model whereby epithelial *Tcf4* Lin- cells represent a stem-cell population that is recruited to, or activated near, sites of colonic barrier damage (Fig. 7A). The *Tcf4* Lin- cells are associated with sequential activation of genes that drive proliferation, followed by those that promote differentiation into multiple colon-cell lineages, to reconstruct the barrier (Fig. 7B). Finally, this epithelial *Tcf4* Lin- stem-cell population, but not the *Tcf4* Lin+ population which contains the Wnt-dependent CBC stem cells, likely represents the cell-of-origin for tumors arising from heterozygous deletion of *Apc* (Fig. 7C).

**Fig. 7:**
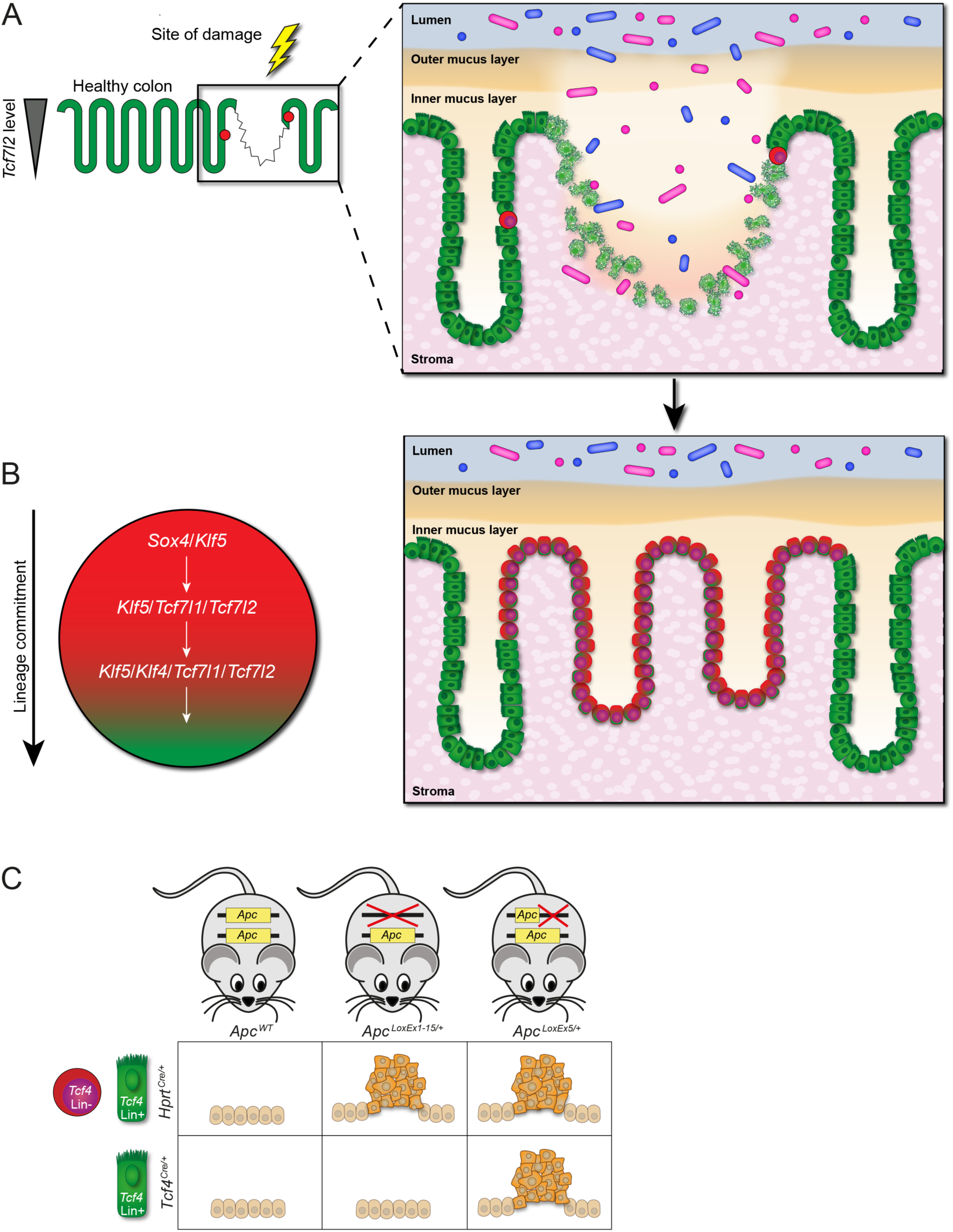
Model depicting the role of Epi *Tcf4* Lin- cells in barrier repair and tumorigenesis. Our data suggest a model whereby (A) Epi *Tcf4* Lin- cells represent a rare pool of stem cells that migrate to and repair sites of damage by replenishing the epithelium and mucus layer with a 1:1 ratio of absorptive to secretory cells, (B) Epi *Tcf4* Lin- stem cells form all cell types of the large intestine upon expression of *Tcf4*, and (C) The Epi *Tcf4* Lin- cell represents the cell-of-origin for colon tumors driven by heterozygous germline deletion of *Apc*.

## Discussion

Several intestinal stem cell populations have been identified including Lgr5+ CBC stem cells, Lrig1+ quiescent stem cells, Krt19+ stem cells, and Dclk1+ stem cells, and these can be distinguished by their unique expression of marker genes, proliferation rate, location and number of cells within the colon crypt (Asfaha et al., 2015; Barker et al., 2007; Chandrakesan et al., 2015; Powell et al., 2012). The Epi *Tcf4* Lin- population therefore represents a rare colonic epithelial stem/progenitor population that is distinct from all previously identified intestinal stem-cell populations. The number and location of stem cells per crypt ranges widely between stem-cell populations. There are approximately 20 Lgr5+ CBC stem cells per crypt (Sasaki et al., 2016), and 1-3 Lrig1+ stem cells per crypt (Powell et al., 2012). Krt19+ stem cells account for 3% of all colon cells (Asfaha et al., 2015), but only 1 in 5 crypts contains a Dclk1+ stem cell (Gagliardi et al., 2012). By comparison, usually a single Epi *Tcf4* Lin- cell is found within only 14% of crypts (Fig. 4Bb-c). Both *Lgr5+* CBC and *Lrig1*+ stem cells exclusively mark cells at the crypt base (Barker et al., 2007; Powell et al., 2012). *Krt19*+ cells are located above the +4 position, up to the isthmus, while Dclk1+ cells are found both at the crypt base and the +4 position (Asfaha et al., 2015; Chandrakesan et al., 2015). We found that Epi *Tcf4* Lin- cells occupy all locations of the crypt but are mostly found in the mid- to upper-crypt region. Cell shape also differs between the stem-cell groups. Lgr5+ CBC stem cells, Lrig1+ stem cells, and Krt19+ cells exhibit a wedge shape (Asfaha et al., 2015), while *Dclk1*+ stem cells have been described with axon-like processes (Gagliardi 2012). In contrast to all other characterized intestinal stem-cell populations, the majority of Epi *Tcf4* Lin- cells are small and round. Intestinal stem cells also differ greatly in proliferative capacity, from a complete lack of proliferation observed in *Dclk1*+ cells (Chandrakesan et al., 2015) to 75% in *Lgr5*+ CBC stem cells (Powell et al., 2012). The rest of the stem-cell populations, including Epi *Tcf4* Lin- cells, show proliferation rates between these two extremes. Finally, previously characterized intestinal stem-cell populations in general have unique gene expression markers. However, *Lgr5*+ CBC and *Lrig1*+ stem cells may have overlapping marker expression (Munoz et al., 2012), raising the possibility that they represent the same stem-cell population. Close comparison of these gene expression profiles demonstrates that only a few markers are shared between Epi *Tcf4* Lin- cells and other identified stem-cell populations. The Epi *Tcf4* Lin- cell therefore represents a distinct, rare colonic epithelial stem/progenitor that is not restricted to a specific location within the colon epithelium.

One interesting aspect of the Epi *Tcf4* Lin- population is that it gives rise to very high numbers of rare epithelial cell types found in the MIX population, such as secretory cells. *Lgr5*+ CBC stem cells are thought to uphold a strict 1:3 secretory-to-absorptive cell ratio, independent of variation in stem-cell numbers (Toth et al., 2017). However, the secretory-to-absorptive ratio in the Epi *Tcf4* Lin- population is 1:1.3 (Tab. S2), and there is a striking enrichment of secretory cells in the MIX population compared to what is found in the normal crypt where absorptive epithelial cells make up >95% of the cells present in the crypt (Fig. 1Ed-e; Colnot et al., 1998). Since secretory cells are critically important during recovery from infection and inflammation following barrier damage (Johansson et al., 2013; Worthington, 2015), we propose that the high concentration of secretory cells derived from the Epi *Tcf4* Lin- population is uniquely suited to the purpose of mucosal recovery. While we have also identified an “absorptive precursor” lineage that is derived from the Epi *Tcf4* Lin-cell, it remains to be determined whether this lineage is independent from absorptive cells arising from the CBC. Interestingly, differentially expressed genes involved in symbiosis, the innate immune response and the antibacterial humoral response are unique to the Epi *Tcf4* Lin- “absorptive precursor” population suggesting that these cells also actively participate in wound repair following injury.

Both Dclk1+ tuft cells and PYY+ enteroendocrine L-cells are commonly associated with intestinal repair (Lebrun et al., 2017; Westphalen et al., 2014). DCLK1+ cells with morphological similarities to tuft cells were observed in the MIX population suggesting that Dclk1 expression is downstream of *Tcf4* expression; however, we cannot rule out the possibility that the Dclk1-expressing cells derived from Epi *Tcf4* Lin- cells do not represent tuft cells but instead are EECs, though morphological characteristics suggest otherwise (Bjerknes et al., 2012). We also identified a colonic cell derived from the Epi *Tcf4* Lin- cell that contains PYY+ granule-filled pseudopod-like neuropods, reminiscent of ileal PYY+ L-cells found in the wound bed periphery following mechanical damage. Since Epi *Tcf4* Lin- cells are rare, it is not surprising that this L-cell type with pseudopod-like processes has not been identified in the colon previously (Bohorquez et al., 2011). Importantly, these cells may interact with enteric glia, which are active mediators of the inflammatory response. In turn, enteric glia may modulate the proliferation and hormone content of the Epi *Tcf4* Lin- PYY+ cells (Bohorquez et al., 2014) to restore mucosal barrier function (Savidge et al., 2007). These findings further support a model whereby Epi *Tcf4* Lin- cells give rise to cells likely to contribute to colonic wound repair.

A role for stem cells in epithelium repair has recently been proposed in which activation of CBCs or de-differentiation of progenitor cells can replenish the gut epithelium in response to modest barrier disruption (Andersson-Rolf et al., 2017). Following proliferation of the epithelium surrounding the wound site, cells may undergo *de novo* crypt formation or crypt fission to complete the wound repair process (Miyoshi et al., 2017; Seno et al., 2009). While the mechanisms that underlie damage repair in the intestine are not well understood, they are thought to include a specialized, migratory WAE repair cell that rapidly seals the wound (Andersson-Rolf et al., 2017; Seno et al., 2009). Importantly, our unique lineage labeling based on *Tcf4* expression has revealed that three days post-injury, EPCAM+ *Tcf4* Lin+ crypts line the surface of the WAE cells in the wound, and an Epi *Tcf4* Lin- and MIX population is apparently expanding from *Tcf4* Lin+ crypts and migrating into the damaged lesion. Our findings therefore suggest that following mechanical damage, an EPCAM+ *Tcf4* Lin+ cell population migrates and contributes to epithelial coverage of the wound bed.

The observation that the Epi *Tcf4* Lin- population appears to expand into injured crypts during wound repair suggests that these cells have the ability to migrate. Indeed, some of the top markers of Epi *Tcf4* Lin- cells are associated with migration, including *Lgals3*, *Agr2*, *Nov*, and *Pigr*. Overexpression of *Lgals3* has been observed in several human cancers and may promote metastatic progression (Yoshimura et al., 2003). *Agr2* has been shown to stimulate proliferation, migration and EMT in salivary adenoid cystic carcinoma (Ma et al., 2017). *Nov* induces migration of renal cell carcinoma cells (Liu et al., 2015). *Pigr* is an inflammation-induced EMT mediator that has also been shown to promote metastatic progression (Ai et al., 2011). We therefore propose a model in which Epi *Tcf4* Lin- stem cells are activated upon injury, migrate to the affected area, and supply the damage site with an abundance of secretory cells and specialized absorptive cells to restore the damaged mucosa.

Our past and present findings suggest that TCF4 functions as a colon tumor suppressor through its role in maintaining the epithelial barrier. We have previously shown that TCF4 is required for the differentiation of CBC stem cells in the colon (Angus-Hill et al., 2011; Korinek et al., 1998). As such, heterozygous deletion of *Tcf4* in the CBC would affect the normal homeostatic process of epithelial barrier renewal. The possibility that TCF4 is important to maintain a functional barrier is supported by increased systemic neutrophil levels in *Tcf4^Het^* animals (Fig. S1). Based on our present findings, reduced levels of TCF4 may also reduce differentiation of the Epi *Tcf4* Lin- population to further inhibit barrier repair. In both stem-cell pathways, reduced TCF4 could promote increased proliferation of stem cells through impairment of differentiation (a cell-autonomous mechanism) or through an increase or persistence of damage signals from unrepaired epithelia (a non-cell-autonomous mechanism), or both. Indeed, we have found that expansion of the *Tcf4* Lin-population in *Tcf4^NeoNull^*animals is likely due to mucosal damage (Angus-Hill et al., 2011). It is important to note that the hypomorphic *Tcf4^CreNeo^* allele may only label descendants of the Epi *Tcf4* Lin- cell in rare damaged crypts and surface epithelium, and not descendants of the CBC. This suggests that the Neo effects may not be at the level of TCF4 expression, as suggested above, but may be related to the effects of Neo on permissive or repressive regulator recruitment to the *Tcf4* promoter in Epi *Tcf4* Lin- and CBC progenitors. Finally, in combination with disruption of the *Apc* gene, barrier dysfunction mediated by reduced TCF4 could explain the synergistic nature of these two gene disruptions during colon tumorigenesis (Angus-Hill et al., 2011). Whether *Apc* loss in these stem-cell pathways affects wound healing remains to be tested.

Interestingly, the Epi *Tcf4* Lin- stem cell population normally lacks expression of both *Tcf4* and *Apc*. When these cells become committed to the secretory and absorptive lineage, *Apc*, *Tcf4*, and *Tcf3* are expressed. The expression pattern of *Apc* in Epi *Tcf4* Lin- cells suggests that it may be important for their lineage commitment, which may explain why heterozygous deletion of the *Apc* gene does not support colon tumorigenesis when introduced specifically in Epi *Tcf4* Lin+ cells. That is, if *Apc* is mutated in the germline (deleted or truncated), lineage commitment might not occur, and the Epi *Tcf4* Lin- stem cell would remain aberrantly proliferative. However, provided normal levels of wild-type *Apc* until the point at which *Tcf4* is expressed, cells with newly-introduced heterozygous deletion of *Apc* may not undergo hyperproliferation since they have completed the early lineage-commitment process. In this model, since heterozygous truncation of *Apc* is able to drive colon tumorigenesis in Epi *Tcf4* Lin+ stem cells, its dominant nature (Schneikert et al., 2007; Tighe et al., 2004) may be sufficient to promote hyperplasia in cells that are more lineage-committed. This hypothesis remains to be tested directly.

Despite evidence to support a role for *TCF4/TCF7L2* as both a tumor suppressor and as a co-oncogene, *TCF4* polymorphisms and mutations are commonly found in human colorectal cancer and other diseases and thereby implicate the loss of *TCF4* function in the disease initiation process (Burwinkel et al., 2006; Grant et al., 2006; Wood et al., 2007). Our studies support this idea and suggest that *Tcf4* functions as a tumor suppressor through its ability to maintain intestinal homeostasis and to stimulate the repair of intestinal barrier damage. Importantly, the ability to isolate and expand normal *TCF4* Lin- cells may ultimately prove useful for the treatment of human patients with diseases caused by intestinal barrier homeostasis defects, such as inflammatory bowel disease (IBD) and irritable bowel syndrome (IBS) (Konig et al., 2016). In addition, experiments designed to understand how mutations in *Tcf4* and *Apc* specifically affect the differentiation of different colon epithelial cell populations could ultimately reveal new genetic mechanisms that drive the development of colon cancer.

## EXPERIMENTAL PROCEDURES

### Mice

The *Hprt^Cre^* mice were obtained and backcrossed into the *C57/BL6* background for over 15 generations (Tang et al., 2002). All other lines have been backcrossed and maintained in the *C57/BL6* background: *Tcf4^Cre^*, *Tcf4^CreNeo^*, *Rosa^mTmG^*, *Apc^LoxEx1-15^* (JAX stock #009045; (Cheung et al., 2010)), *Apc^LoxEx5^*(KOMP; APC tm1c; Reporter removed with FlpE recombination (Fig. 6B), as animals with reporter have a phenotype in the absence of Cre, and no phenotype without reporter until conditional deletion), *Cd2^iCre^* (JAX stock #008520; (de Boer et al., 2003; Shimshek et al., 2002)). Mice were injected i.p. with 50 mg/kg of EdU either two hours (FACS) or 16 hours (Crypt Isolation) prior to sacrifice. Studies and procedures performed on mice strictly followed the animal protocol approved by the University of Utah Institutional Animal Care and Use committee (IACUC).

### Statistics

Throughout, data from at least 3 independent experiments were analyzed and statistics were expressed as mean ± SEM. P-values were calculated using a two-tailed Student’s *t*-test with statistical significance P < 0.05 (*), P < 0.005 (**), or P < 0.0005 (***).

### Genotyping

DNA was isolated from animal tail biopsies using the HOTShot system: tissues were heated to 95°C for 60 minutes in 75 µL of alkaline lysis reagent (25 mM NaOH, 0.2 mM Disodium EDTA adjusted to pH 12), and neutralized in neutralizing reagent (40 mM Tris-HCl adjusted to pH 5). PCR was performed using 50 ng of the prepared template DNA solution and a final concentration of 0.25 µM of each primer. The primer pairs Ex5-F (5’ - ATACCGTACATTGCTTGGGGAAAGG - 3’) and Ex5-R (5’ - GCAAGCAGATAGTATAATGTTTCTACCCAT - 3’) were used to detect the wild type, floxed, and recombined alleles for the APCEx5 variant. The primer pairs Ex15-F (5’ – CCTGCCCCCTCTTATTTTCT −3’) and Ex15-R (5’ – CACGAGGATAATGGCCAAAC – 3’) were used to detect the wild type and floxed alleles, and Ex15-F and Ex15-R2 (5’ – GGCTGGGTAAACATATTCATGGG – 3’) were used to detect the recombined allele for the APCex15 variant.

### Crypt isolation and tissue dissociation

To isolate crypts for single cell preparations, we adapted a procedure that separates colonic epithelium from the underlying mesenchyme (Nik and Carlsson, 2013). A gavage needle with silicone tip attached to a 3 ml syringe was inserted into the cleaned colon from the distal end and pulled all the way through to the proximal end. The plunger needed to be pulled out and the barrel filled with air. The proximal part of the colon was secured to the tip of the gavage needle with thread. Using a thumb, the colon was pulled over the tip of the needle, thus inverting it. Following inversion, the proximal end of the colon was ligated with thread while the other end was still fitted to the gavage needle. The inverted colon was then submerged in 3.8 mL of ice-cold Cell Recovery Solution (Corning) in a 5 mL round-bottom polystyrene tube (Corning) and placed on ice. The submerged colon was inflated and deflated every 5 minutes with occasional swirling in the solution for 1 hour. During this time, the Cell Recovery Solution dissolved the basement membrane and the air pressure pushed the crypts out of the crypt beds. A 10-minute digestion with TrypLE^TM^ Express (Thermo Fisher) at 37°C then generated a single cell suspension. Addition of FBS and wash buffer (DPBS containing 3% BSA) inhibited the TrypLE^TM^ digest. The single cell suspension was filtered using a 100 µm cell strainer and washed. Wash buffer used to prepare cells for Drop-seq procedure contained 0.5 mM EDTA.

To generate a whole crypt suspension, the colon was cut into small pieces after incubation in Cell Recovery Solution and inflation and deflation steps (or immediately following dissection). Colon pieces were placed into an EDTA Chelation buffer for 40 minutes while shaking at 37°C. Crypts were then fixed in 4% PFA and washed using PBS. The crypt-containing supernatant was harvested for further staining and imaging.

### Crypt counts

Crypt suspensions were used to count the number *Cd2* Lin+ cells (in the *Cd2^iCre^ Rosa^mTmG^* mouse model) or *Cd2* Lin- and *Tcf4* Lin- cells (in the *Tcf4^Cre^ Cd2^iCre^ Rosa^mTmG^* mouse model) in 100 randomly selected crypts via confocal analysis. The location within the crypt was noted (crypt base, +4 position, mid, or top crypt).

### Mouse Tissue Immunofluorescence and Immunohistochemistry

Crypt suspensions were prepared as above described. Frozen tissue sections were prepared and stained using conventional methods. Primary antibodies were anti-DCLK1 (Novus Biologicals), anti-HES1 (Abcam), anti-SOX9 (Abcam), anti-IHH (R&D Systems), anti-PYY (Abcam), and AlexaFluor^®^ 647-conjugated anti-DLL1 (BioLegend). Antibodies against the antigens DCLK1, HES1 and IHH were conjugated to allophycocyanin (APC) using a Lightning-Link APC Antibody Labeling Kit (Novus Biologicals). The secondary antibody used in combination with anti-PYY antibody was an AlexaFluor^®^ 647 conjugate (Abcam). DAPI (Thermo Fisher) was utilized as DNA stain. Confocal imaging for crypt suspensions, surface epithelium, frozen tissue sections, crypt counts, and single cells collected via flow cytometry was performed with an Olympus FV1200 confocal microscope attached to an Olympus IX81 Inverted Microscope.

### Flow Cytometry

Single cells obtained from crypt isolation and subsequent tissue dissociation were stained with AlexaFluor^®^ 647-conjugated anti-CD8a (BioLegend) antibody for 1 hour on ice. Alternatively, the Click-iT^®^ Plus EdU AlexaFluor^®^ 647 Flow Cytometry Assay Kit (Thermo Fisher) was used according to the manufacturer’s instructions for the labeling of EdU (Animals collected 2 hours post-EdU injection). Hoechst 33324 (Thermo Fisher) was used as a DNA stains. Staining with CellTrace^TM^ Calcein violet AM (Thermo Fisher) according to the manufacturer’s instructions was for distinguishing live cells. The University of Utah Flow Cytometry Facility performed cell sorting with a 5-laser 13-detector BD FACSAria Cell Sorter (BD Biosciences). We eliminated dead cell by gating forward/side scatter and selecting TOM and/or GFP positive cells. Sorted cells were collected in 1.5 mL microcentrifuge tubes containing either wash buffer (DPBS containing 3% BSA) for subsequent confocal analysis or Complete DMEM + EDTA for Drop-seq. Data analysis was performed with FlowJo software.

### Drop-seq procedure

Isolated Tcf4 Lin- and Tcf4 MIX populations were separately combined with human cells to reach optimal cell number and concentration for Drop-seq, and single-cell suspensions were diluted to 270 cells/ml and processed as described previously (Macosko et al., 2015). Briefly, cells, barcoded microparticle beads (ChemGenes Corporation, Lot #113015B), and lysis buffer were co-flown into a microfluidic device and captured in nanoliter sized droplets. After droplet collection and breakage, the beads were washed, followed by cDNA synthesis on the bead using Maxima H-minus RT (Cat # Thermo Fisher Scientific). Excess oligos were removed via exonuclease I digestion. cDNA amplification was done from a pool of 2,000 beads using HotStart ReadyMix (Kapa Biosystems). Individual PCRs were purified and pooled for library generation. A total of 600pg of amplified cDNA was used for a Nextera XT library preparation (Illumina), and the New-P5-SMART PCR hybrid oligo, and a modified P7 Nextera oligo with 10 bp barcodes was used for sequencing. A 125 cycle paired-end sequence run was performed on an Illumina HiSeq 2500 instrument using HiSeq SBS kit v4 sequencing reagents and the Read1CustomSeqB primer as described (Macosko et al., 2015).

### R-based analysis of Drop-seq data

Drop-seq data was processed with the McCarroll pipeline (Macosko et al., 2015). 1117 mouse cells were sequenced and single cell data analysis was accomplished using the Seurat package (Script S1-S4; Seurat version 2.0.1; http://satijalab.org/seurat/). Following several filtration steps, a total number of 1,023,641 counts were identified with an average of 324 genes expressed per cell (>1count). In total, 381 unique genes were expressed across 516 cells, and principal component analysis (PCA) was used to identify statistically significant PCs. From a plot of the standard deviation of PCs, the first five PCs were selected (Fig. S6A). Based on the Euclidean distance in PCA space, a K-nearest neighbor (KNN) graph was constructed and modularity optimization techniques were applied to cluster cells. The dataset was visualized using t-Distributed Stochastic Neighbor Embedding (tSNE) (Fig. 2A,C) and the most differentially expressed genes for each cluster were calculated (Tab. S1; Tab. S3).

### Monocle

To specifically select for and analyze Epithelial Pigr+ cells, we filtered for cells with *Pigr* expression and the absence of T cell markers and selected genes with high dispersion and expression. Monocle then constructed single cell trajectories, featuring three cell states and one branch point. Pseudotime was plotted and helped visualize one cell state being the starting point and then branching off into two different states as time passes (Fig. 3A, Script S5-6).

### Endoscopic Damage

Food was withheld from mice the afternoon before surgery. In preparation of the colonoscopy the following morning, mice were anesthetized using a system for isoflurane anesthesia (VetEquip) that allows simultaneous flow of isoflurane and oxygen. A mouse colonoscope (Karl Storz) was inserted into the animal’s rectum and insufflation was used to inflate the colon during the procedure. Flexible biopsy forceps with a diameter of 3 French were inserted into the working channel of the colonoscope and six biopsy samples were taken. The mice were closely monitored during procedure recovery. Three (or five) days post-surgery the mice were sacrificed and colonic damage sites inspected using a dissecting microscope. Smaller wounds were cut out with a 2 mm Sterile Disposable Biopsy Punch (Miltex), larger wounds with a 4-mm biopsy punch. To prevent curling of biopsy punches, cover slips weighed down the punches during paraformaldehyde fixation. Then biopsy punches were frozen, prepared, and stained using conventional methods.

### Mouse Tissue Immunofluorescence and Immunohistochemistry

Crypt suspensions were prepared as above described. Frozen tissue sections were prepared and stained using conventional methods. Primary antibodies were anti-DCLK1 (Novus Biologicals), anti-HES1 (Abcam), anti-SOX9 (Abcam), anti-IHH (R&D Systems), anti-PYY (Abcam), anti-EPCAM (Invitrogen), and AlexaFluor^®^ 647-conjugated anti-DLL1 (BioLegend). Antibodies against the antigens DCLK1, HES1 and IHH were conjugated to allophycocyanin (APC) using a Lightning-Link APC Antibody Labeling Kit (Novus Biologicals). The secondary antibody used in combination with anti-PYY and anti-EPCAM antibodies was an AlexaFluor^®^ 647 conjugate (Abcam). The Click-iT^®^ Plus EdU AlexaFluor^®^ 647 Flow Cytometry Assay Kit (Thermo Fisher) was used to stain EdU-marked proliferating cells (animals were injected with EdU two hours prior to sacrifice) on biopsy punches. DAPI (Thermo Fisher) was utilized as DNA stain. Confocal imaging for crypt suspensions, surface epithelium, frozen tissue sections, crypt counts, and single cells collected via flow cytometry was performed with an Olympus FV1200 confocal microscope attached to an Olympus IX81 Inverted Microscope.

### ACCESSION NUMBERS

Single-cell sequencing data for colonic Tcf4 Lin- cells are publicly available through the Gene Expression Omnibus (GEO) under accession number **GSE129517**.

## SUPPLEMENTAL INFORMATION

Figure S1: Neutrophils

Figure S2: Edu Staining of Tcf4GFPCre

Figure S3: Hes1 staining

Figure S4: Feature Plots B cells + Epi+

Figure S5: Feature Plots T cells

Figure S6: PC analysis

Figure S7: Trajectory

Figure S8: Cd2 cells

Figure S9: Apc

Table S1: Diff. expression Tcf4Lin- cells

Table S2: Seurat cluster assignments

Table S3: Diff. expression Tcf4Lin- Epi cells

Table S4: Cd2iCre crypt counts

Table S5: Single cell seq data

Table S6: Cd2iCre tumor count

Supplemental Video 1: Colonoscopy Damage

Script S1: Seurat_Tcf4Lin- cell.Rmd

Script S2: Seurat_Tcf4Lin- cell.html

Script S3: Seurat_Tcf4Lin- Epi cell.Rmd

Script S4: Seurat_Tcf4Lin- Epi cell.html

Script S5: Monocle_Tcf4Lin- cell.Rmd

Script S6: Monocle_Tcf4Lin- cell.html

## Supporting information

All_Supplemental_Figures_and_Legends

SuppTable_Description_ReadMe

Table_S5_Single_cell_seq_data__01

Table_S6_Cd2iCre_tumor_count__01

Table_S1_Diff._expression_Tcf4Lin-_cells__01

Table_S1_Diff._expression_Tcf4Lin-_cells__02

Table_S1_Diff._expression_Tcf4Lin-_cells__03_MAH_X1_X2_resultsC1

Table_S1_Diff._expression_Tcf4Lin-_cells__04_MAH_X1_X2_resultsC2

Table_S1_Diff._expression_Tcf4Lin-_cells__05_MAH_X1_X2_resultsC3

Table_S2_Seurat_cluster_assignments__01_cluster_assignments_ALL

Table_S2_Seurat_cluster_assignments__02_cluster_assignments_Epi_cell

Table_S3_Diff._expression_Tcf4Lin-_Epi_cells__01

Table_S3_Diff._expression_Tcf4Lin-_Epi_cells__02

Table_S3_Diff._expression_Tcf4Lin-_Epi_cells__03

Table_S3_Diff._expression_Tcf4Lin-_Epi_cells__04

Table_S4_Cd2iCre_crypt_counts__01

Table_S4_Cd2iCre_crypt_counts__02

Table_S4_Cd2iCre_crypt_counts__03

Table_S4_Cd2iCre_crypt_counts__04

Table_S4_Cd2iCre_crypt_counts__05

Colonoscopy damage video

## ACKNOWLEDGMENTS

We would like to thank Cicely Jette for editing, critical reading, and helpful discussions. Special thanks to Diana L. Lim for original artwork and figure formatting throughout the manuscript, and for critical reading by A. Boulet. We acknowledge Z. Ren for histological tumor analysis, and technical assistance from C. Brown, and J. Marvin in the Flow Cytometry core. We acknowledge KOMP, C. Lenz, S. Barnett, K. Lustig and the Capecchi laboratory for providing the *Apc^LoxEx5^* cell line, for cell-line injections and chimera production. The research reported in this publication was supported by access to technical cores supported by National Cancer Institute award P30 CA042014. The project described was supported by the National Cancer Institute Grants P01 CA073992 and K01 CA128891 and the Huntsman Cancer Foundation.

## Notes

https://www.ncbi.nlm.nih.gov/geo/query/acc.cgi

### Competing Interest Statement

The authors have declared no competing interest.

