## Supplementary material for "A Rare T-Cell Factor 4 Lineage-negative Epithelial Stem Cell Supports Wound Repair and APC-deletion-induced Colon Tumorigenesis": All_Supplemental_Figures_and_Legends

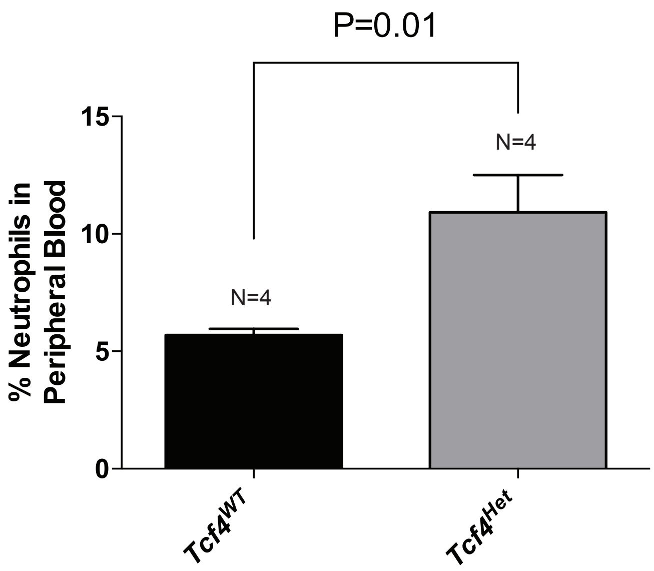


**Fig. S1: Percentage of Peripheral Neutrophils is higher in *Tcf4$^{Het}$* animals compared to *Tcf4$^{WT}$* males, related to Fig.1.** P=<0.05 using two-sample t-test with welch correction. Error bars were calculated using SEM.

A

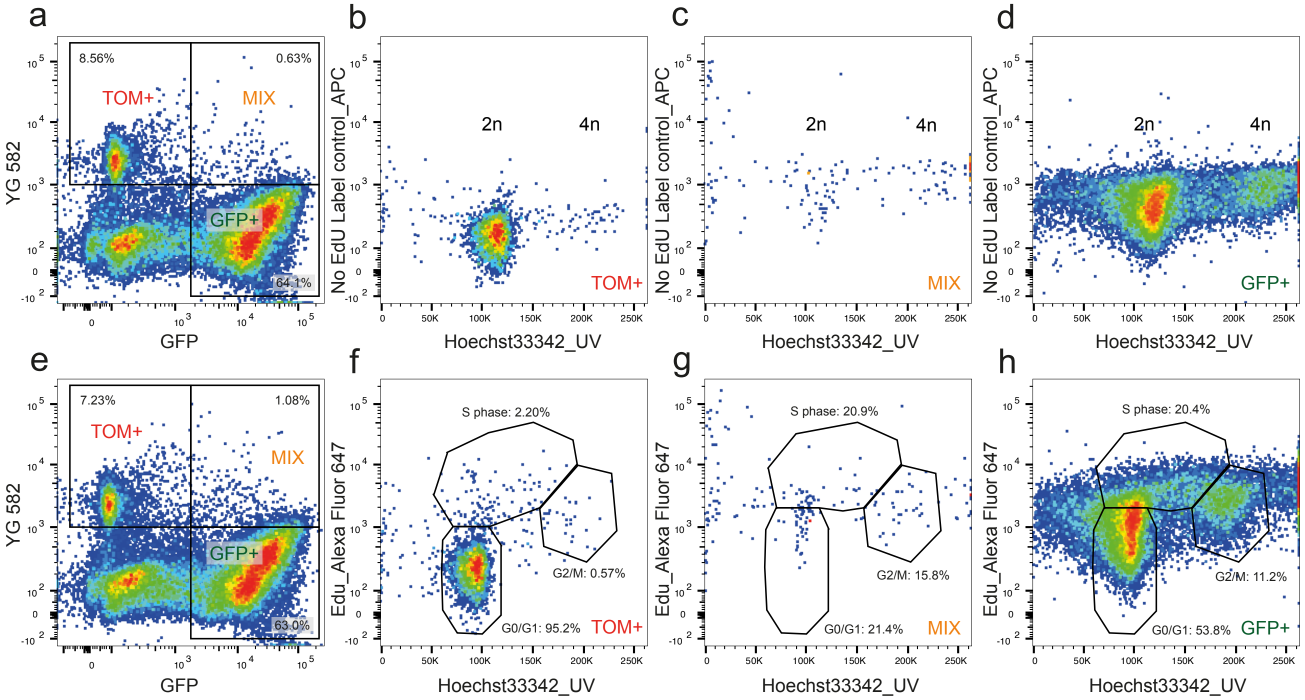


B

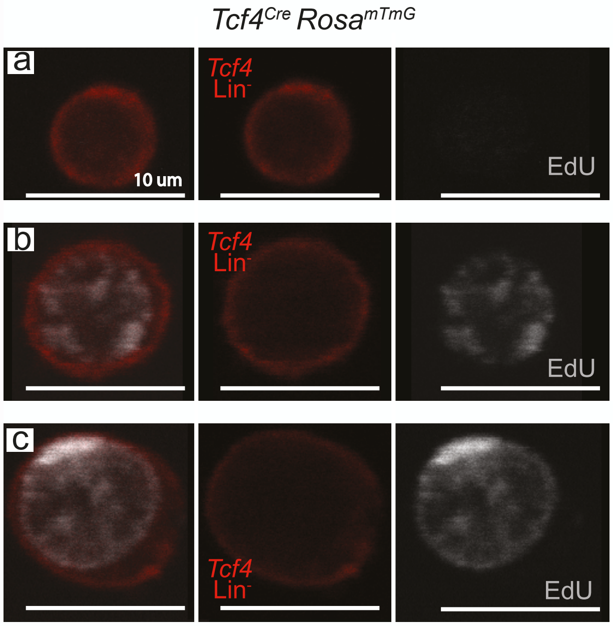


**Fig. S2: Proliferation and cell cycle studies *Tcf4^Cre Rosa^mTmG* mouse, related to Fig. 1.** (A) Representative flow cytometric analysis of (Aa-d) unlabeled and (Ae-h) cell cycle analysis of the EdU labeled populations. (B) Characterization of FACs sorted Tom+ Edu+ cells by confocal microscopy.

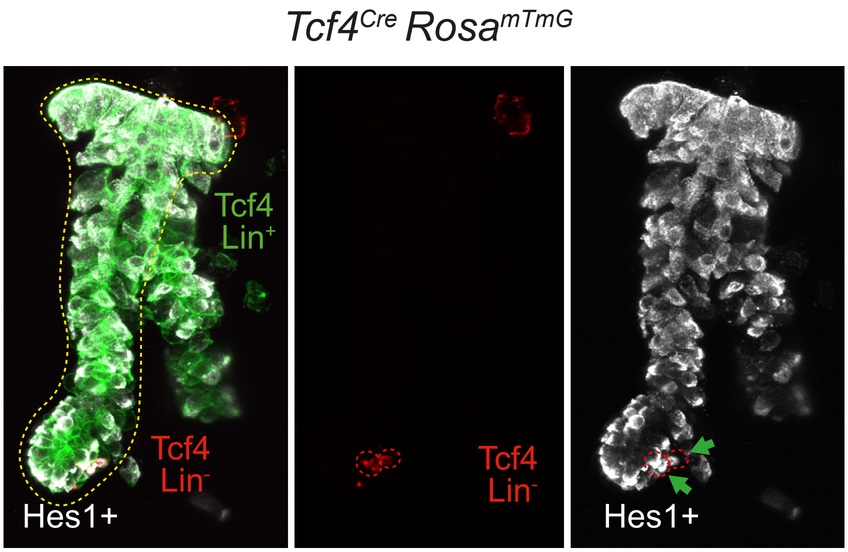


**Fig. S3: Co-staining of the Tcf4 Lin- population and HES1, related to Fig. 1**. Immunofluorescence staining of colonic crypts of a *Tcf4*$^{Cre}$ *Rosa*$^{mTmG}$ mouse with anti-Hes1 antibody. Dotted red lines encircle *Tcf4* Lin- cells. Green arrows denote *Tcf4* Lin- cells that express the HES1.

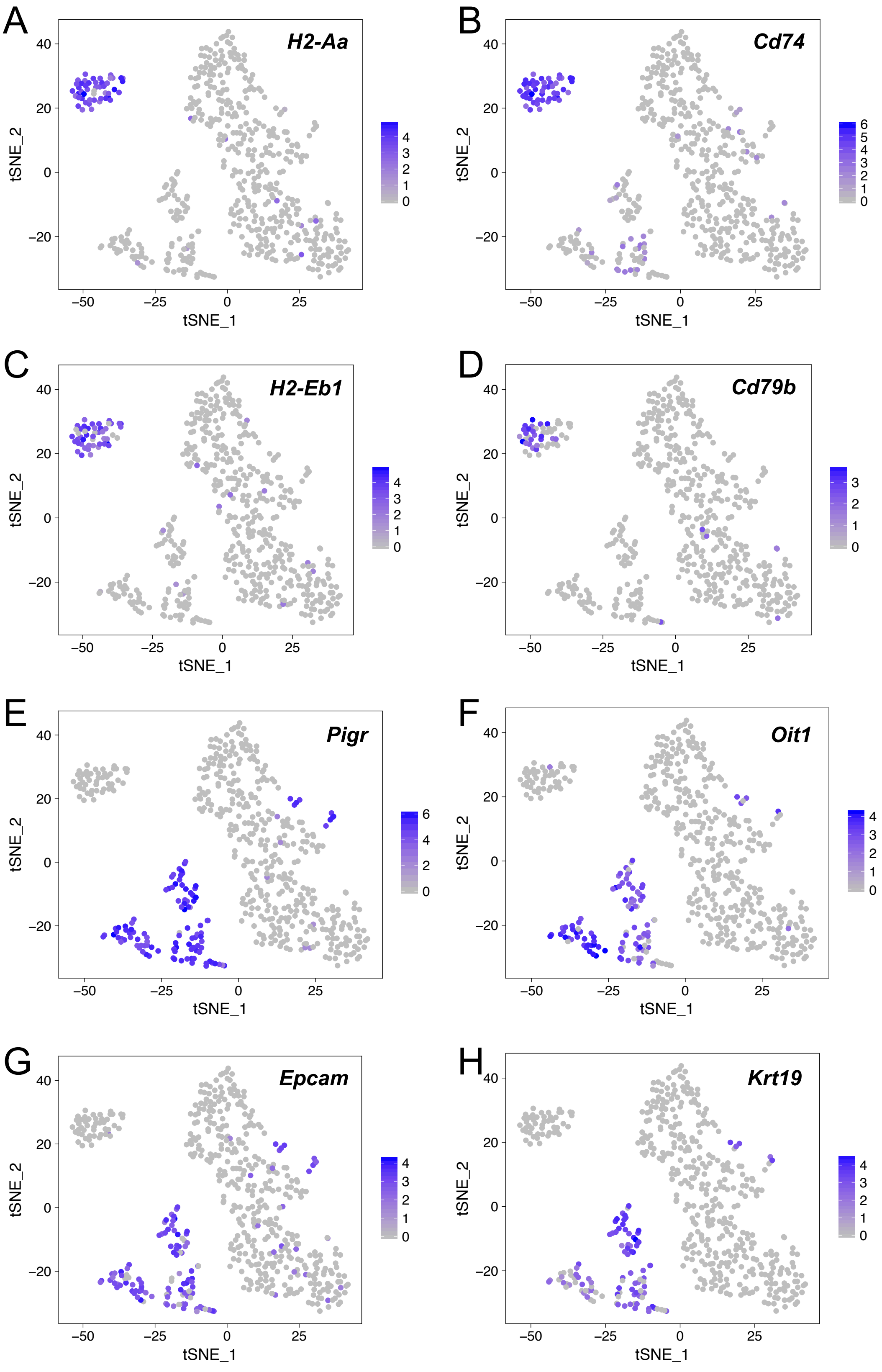


**Fig. S4**: **Expression of B-cell and epithelial cell markers in *Tcf4* Lin- cells, related to Fig. 2.** Feature Plots are a tool in the R package Seurat and visualize marker expression, including expression levels, on a tSNE plot. (A-D) Feature Plots of B-cell markers. (A) *H2-Aa*. (B) *Cd74*. (C) *H2-Eb1*. (D) *Cd79b*. (E-H) Feature Plots of epithelial cell markers. (E) *Pigr*. (F) *Oit1*. (G) *Epcam*. (H) *Krt19*.

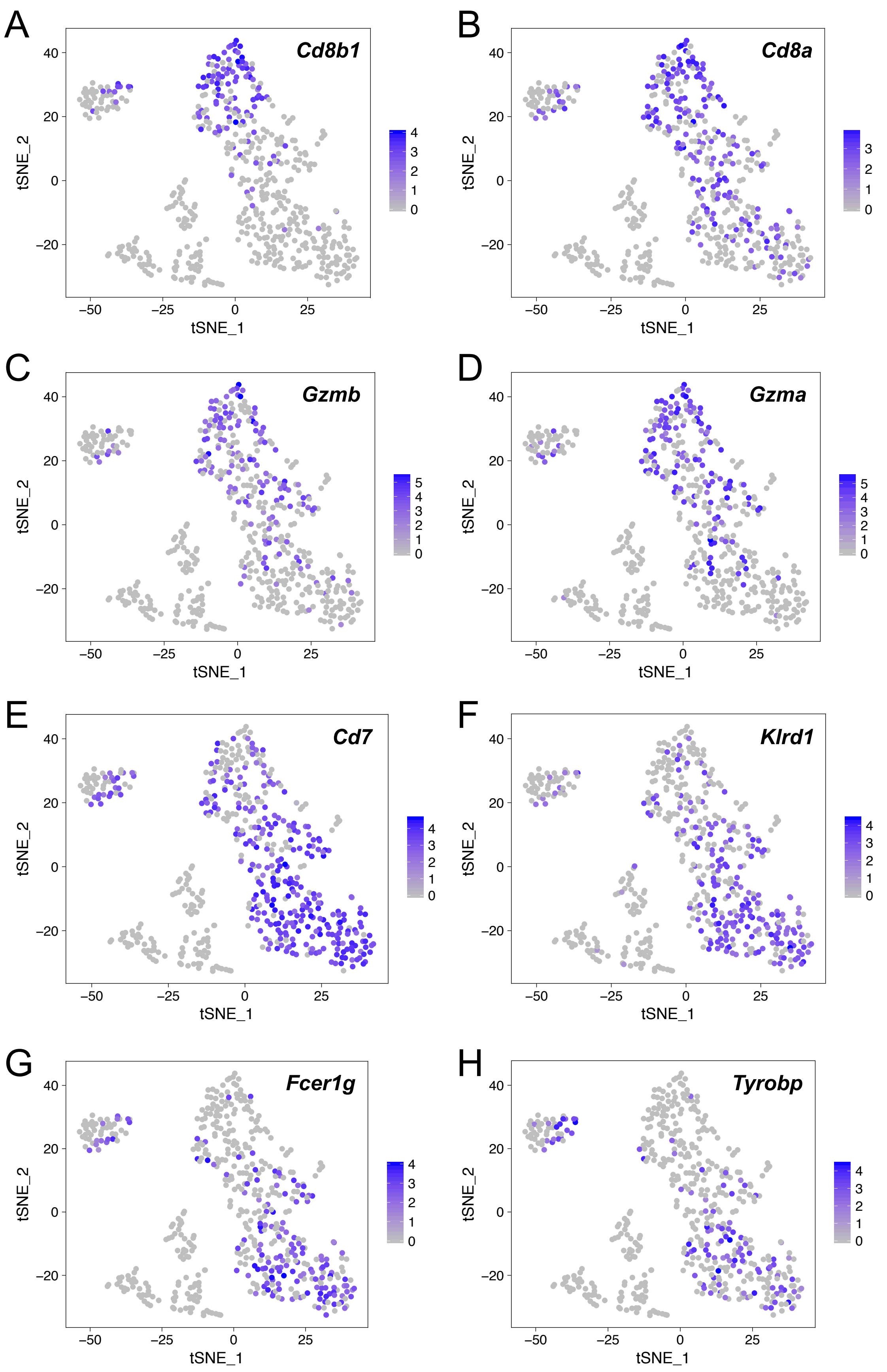


**Fig. S5: Expression of IEL markers in *Tcf4* Lin- cells, related to Fig. 2.** Feature Plots are a tool in the R package Seurat and visualize marker expression, including expression levels, on a tSNE plot. (A-D) Feature Plots of iIEL markers. (A) *Cd8b1*. (B) *Cd8a*. (C) *Gzmb*. (D) *Gzma*. (E-H) Feature Plots of nIEL markers. (E) *Cd7*. (F) *Klrd1*. (G) *Fcer1g*. (H) *Tyrobp*.

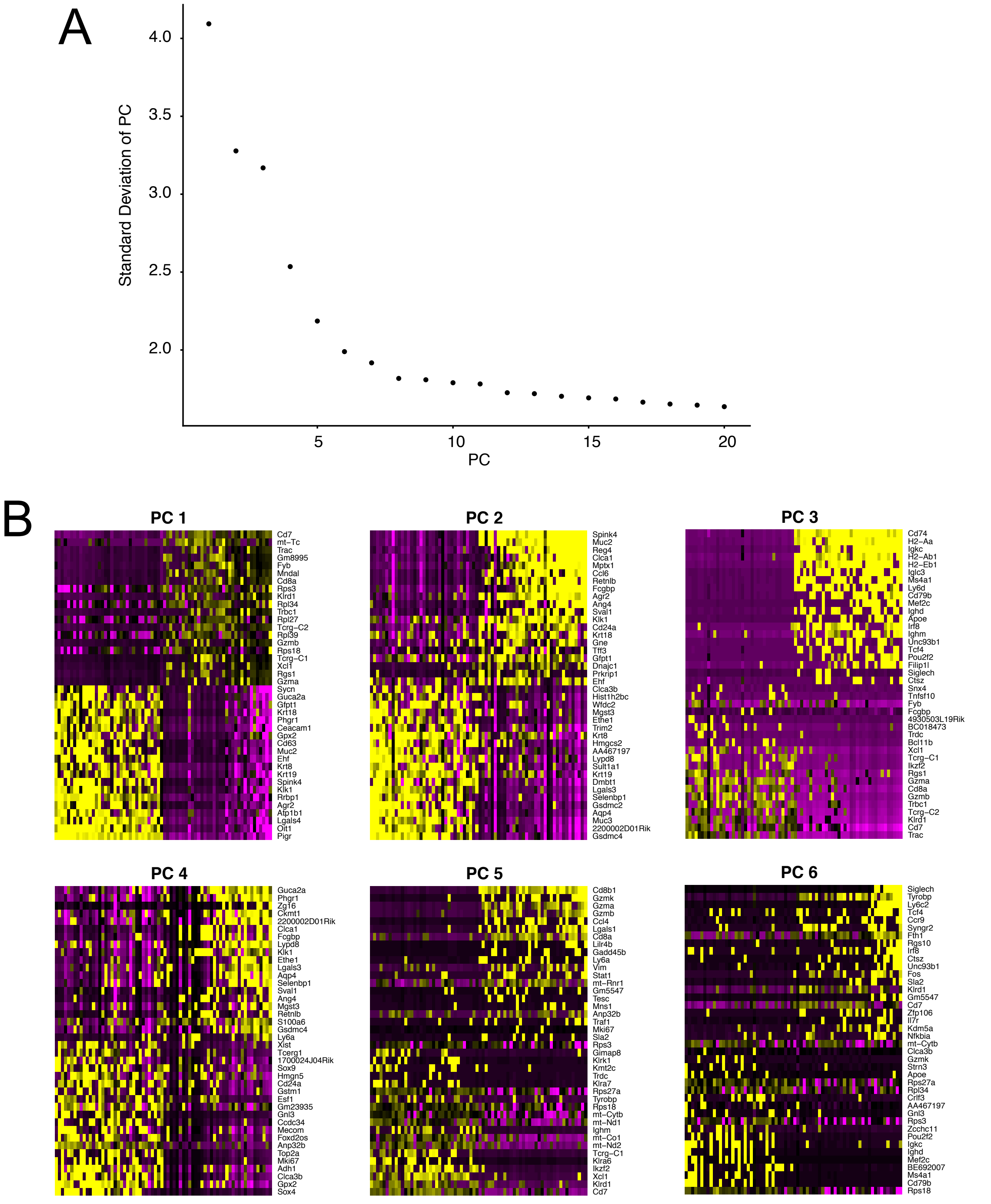


**Fig. S6**: **Identification of statistically significant principal components, related to Fig. 2.** Analysis of single cell RNA-sequencing data utilizing the R package Seurat. (A) Plot of the standard deviations of principle components to determine which principle components to include in further analysis. (B) PCHeatmap displays heterogeneity of dataset by ordering cells and genes according to their principal component analysis (PCA) scores.

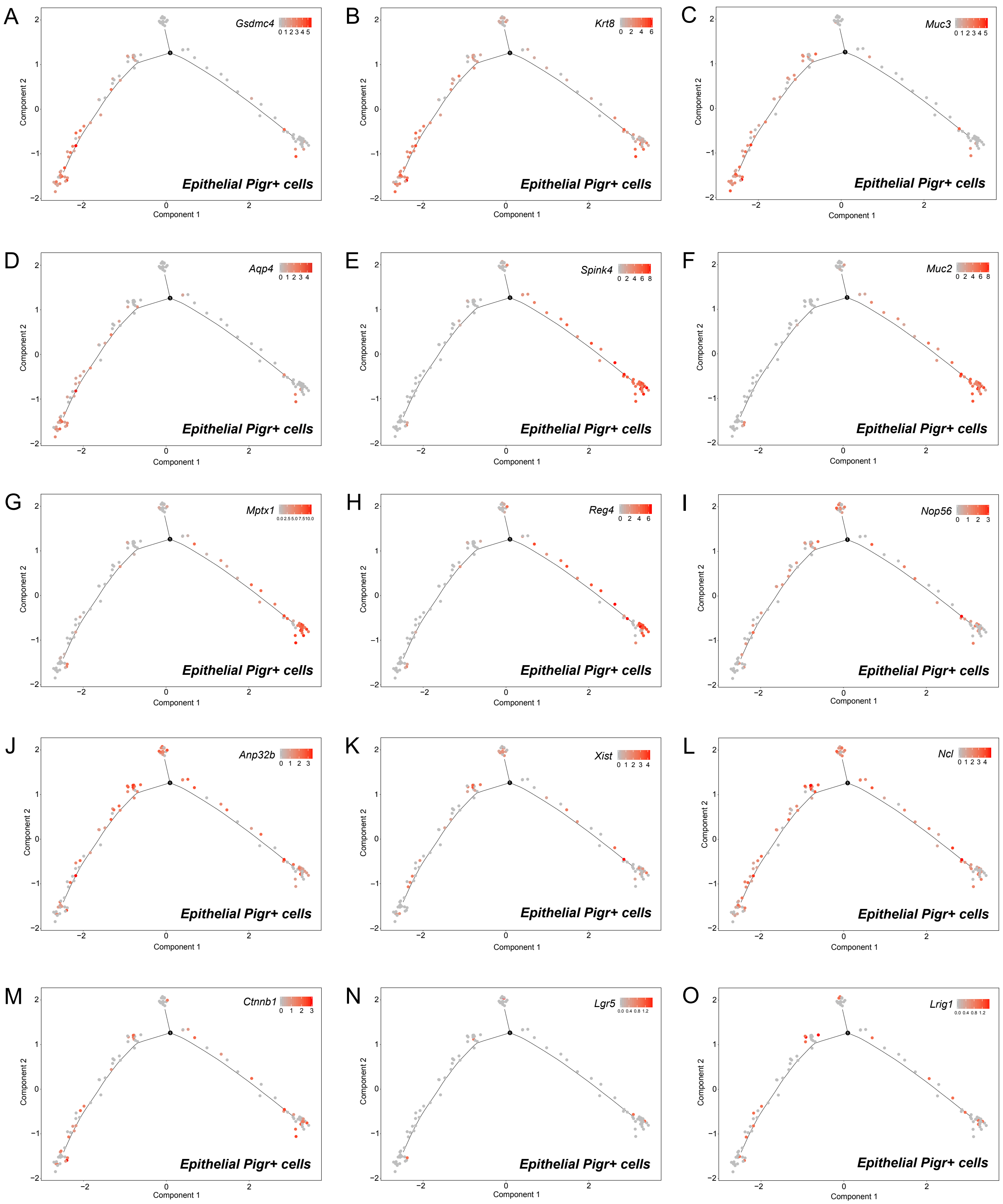


**Fig. S7: Organization of Epi *Tcf4* Lin- cells in pseudotime confirms that a stem cell cluster gives to alternative cell fates, either absorptive or secretory, related to Fig. 3.** Analysis of the pseudotime properties of Epi *Tcf4* Lin- cells by employing the monocle algorithm. (A-O) Single cell trajectories displaying the expression level of a marker and placing cells in pseudotime. (A) *Gsdmc4*. (B) *Krt8*. (C) *Muc3*. (D) *Aqp4*. (E) *Spink4*. (F) *Muc2*. (G) *Mptx1*. (H) *Reg4*. (I) *Nop56*. (J) *Anp32b*. (K) *Xist*. (L) *Ncl*. (M) *Ctnnb1*. (N) *Lgr5*. (O) *Lrig1*.

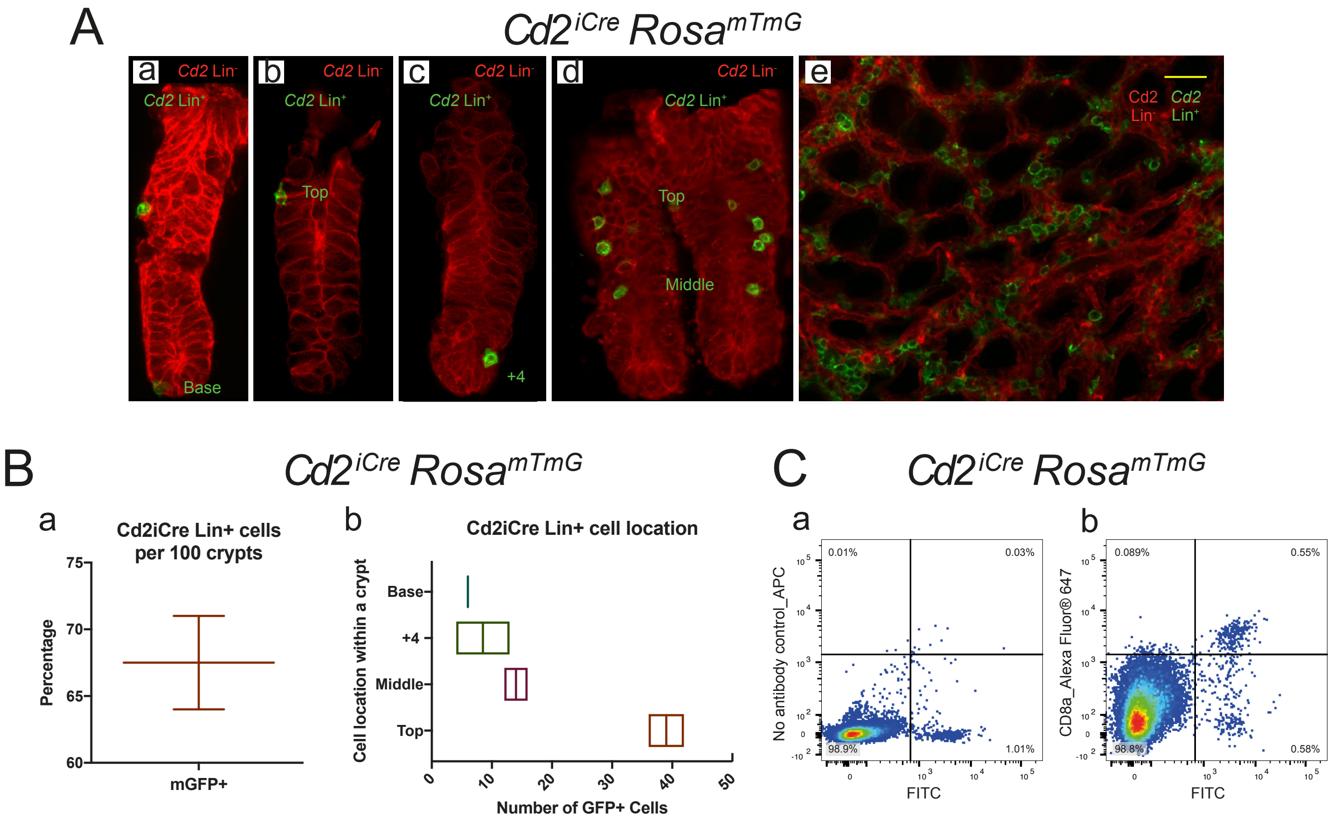


**Fig. S8**: **Lineage analysis of *Cd2*$^{iCre}$ expression in colonic crypts of *Cd2*$^{iCre}$ *Rosa*$^{mTmG}$ mouse, related to Fig. 4.** (Aa-d) Localization of *Cd2*$^{iCre}$ positive cells (mGFP) throughout the crypt and (Ae) in the intercryptal region of the colon surface following crypt isolation. (B) mGFP+ cell numbers per 100 crypts and crypt location of *Cd2*$^{iCre}$ Lin+ cells. (C) Representative FACs of anti-Cd8a stained single colon crypt cells isolated from *Cd2*$^{iCre}$ lineage. (scale bar, 50 μm).

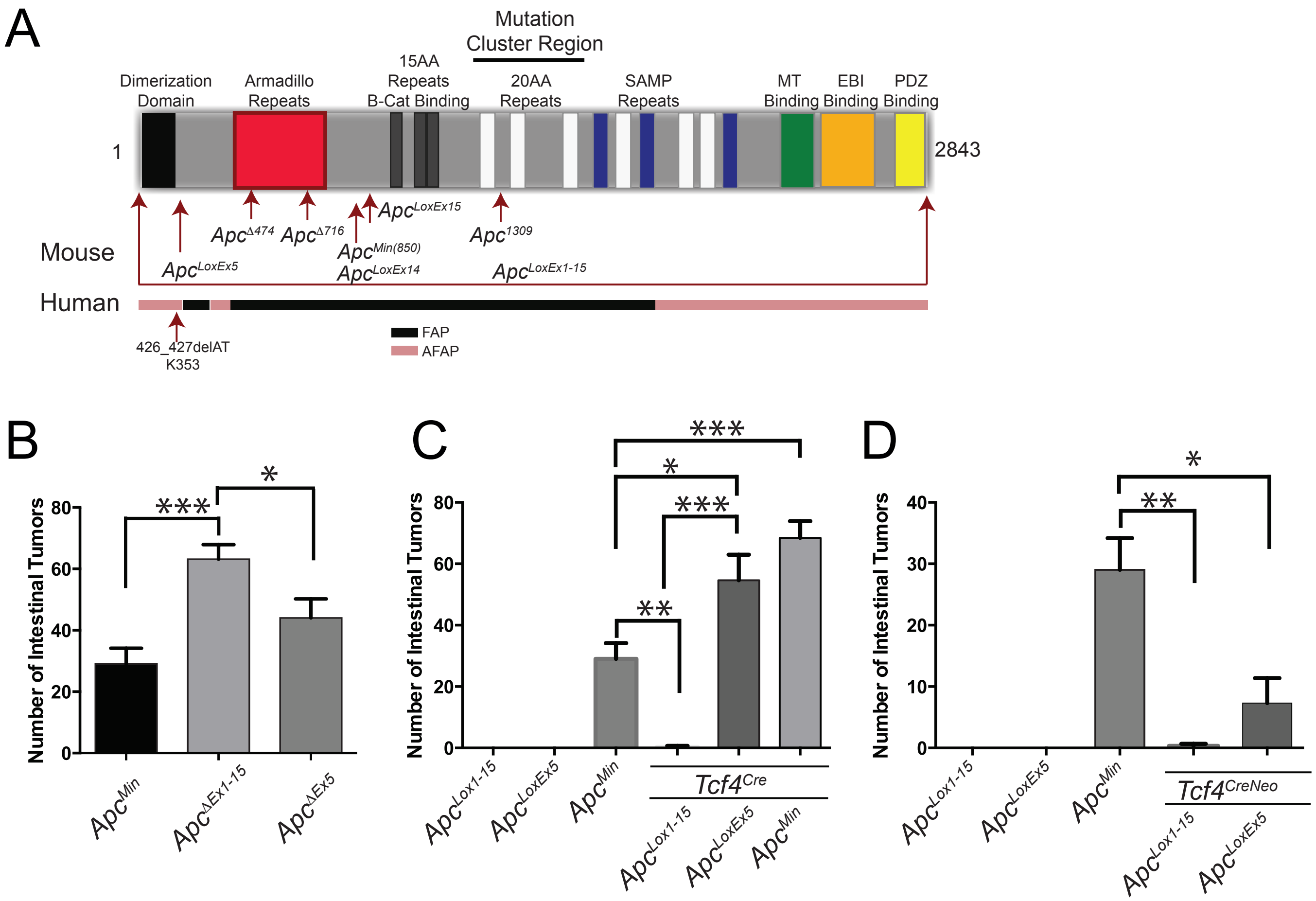


**Fig. S9**: **Cartoon depicting location of *Apc* mutations in common mouse models and Cre-driven intestinal tumorigenesis, related to Fig. 6.** Intestinal tumorigenesis following recombination of (A) *Apc*$^{LoxEx1\text{-}15}$ and *Apc*$^{LoxEx5}$ in (B) the germline using *Hprt*$^{Cre}$, (C) *Tcf4*$^{Cre}$ and (D) *Tcf4*$^{CreNeo}$ driven recombination. Intestinal tumorigenesis counts as described in Fig. 6 of the main text. *p = 0.05, **p = 0.005, ***p = 0.0005 using two–sample t- test with Welch correction. Error bars were calculated using SEM.
