## Supplementary material for "A Rare T-Cell Factor 4 Lineage-negative Epithelial Stem Cell Supports Wound Repair and APC-deletion-induced Colon Tumorigenesis": SuppTable_Description_ReadMe

**Supplemental Tables description:**

**Table S1: Diff. expression Tcf4Lin- cells**

Description: For all Tcf4Lin- cells, genes expressed in each individual cluster were compared to all other cells and a list with most differentially expressed genes printed. The list shows genes that can be detected at a minimum percentage of 25%.

Worksheet exports:

- Table_S1_Diff._expression_Tcf4Lin-_cells__01_Summary.csv

- Table_S1_Diff._expression_Tcf4Lin-_cells__02_MAH_X1_X2_resultsC0_25percent.t.csv

- Table_S1_Diff._expression_Tcf4Lin-_cells__03_MAH_X1_X2_resultsC1_25percent.t.csv

- Table_S1_Diff._expression_Tcf4Lin-_cells__04_MAH_X1_X2_resultsC2_25percent.t.csv

- Table_S1_Diff._expression_Tcf4Lin-_cells__05_MAH_X1_X2_resultsC3_25percent.t.csv

**Table S2: Seurat cluster assignments**

Description: Cluster assignment of each individual cell in the tSNE analysis of Seurat.

Worksheet exports:

- Table_S2_Seurat_cluster_assignments__01_cluster_assignments_ALL.csv

- Table_S2_Seurat_cluster_assignments__02_cluster_assignments_Epi_cell.ts.csv

**Table S3: Diff. expression Tcf4Lin- Epi cells**

Description: For all Tcf4Lin- Epi cells, genes expressed in each individual cluster were compared to all other cells and a list with most differentially expressed genes printed. The list shows genes that can be detected at a minimum percentage of 25%.

Worksheet exports:

- Table_S3_Diff._expression_Tcf4Lin-_Epi_cells__01_Summary.csv

- Table_S3_Diff._expression_Tcf4Lin-_Epi_cells__02_MAH_X1_X2_resultsC0_PIGR_25perc.csv

- Table_S3_Diff._expression_Tcf4Lin-_Epi_cells__03_MAH_X1_X2_resultsC1_PIGR_25perc.csv

- Table_S3_Diff._expression_Tcf4Lin-_Epi_cells__04_MAH_X1_X2_resultsC2_PIGR_25perc.csv

**Table S4: Cd2iCre crypt counts**

Description: The number and location of green cells in isolated crypts of *Cd2iCre* *Rosa^mTmG^* mice and red cells in isolated crypts of *Tcf4^Cre^ Cd2^iCre^ Rosa^mTmG^* mice. 100 crypts were counted per colon.

Worksheet exports:

- Table_S4_Cd2iCre_crypt_counts__01_950_10.csv

- Table_S4_Cd2iCre_crypt_counts__02_954_2.csv

- Table_S4_Cd2iCre_crypt_counts__03_951_1.csv

- Table_S4_Cd2iCre_crypt_counts__04_950_8.csv

- Table_S4_Cd2iCre_crypt_counts__05_955_2.csv

**Table S5: Single cell seq data**

Description: Single cell gene read per barcode.

Worksheet exports:

- Table_S5_Single_cell_seq_data__01_MAH_X1_X2_counts.txt.csv

**Table S6: Cd2iCre tumor count**

Description: Tumor counts for several *ApcLox^Ex1-15^ Cd2^iCre^* or *ApcLox^Ex5^ Cd2^iCre^* mice.

Worksheet exports:

- Table_S6_Cd2iCre_tumor_count__01_Tumor_count.csv
